# The progression of sex differences in brain networks across the lifespan

**DOI:** 10.64898/2026.01.30.702608

**Authors:** Ke Huang, Keith W. Jamison, Emily Jacobs, Nina Miolane, Amy Kuceyeski

## Abstract

Sex differences in brain connectivity are well documented, yet how these differences evolve across the human lifespan remains poorly understood. Rigorously assessing sex-dependent trajectories of brain network organization is challenging due to difficulty in acquiring, processing, and modeling high-dimensional connectomes. Here, we analyzed 15 types of functional and structural connectivity networks from 1286 healthy individuals aged 8–100+ years, using our new AI-based Krakencoder to derive a low-dimensional multimodal “fusion” connectome representation. Sex differences were minimal in early childhood, pronounced in young to mid-adulthood, and diverged across modalities in later life: functional connectivity grew less distinct and structural connectivity grew more distinct from midlife onward. Functional differences were driven predominantly by higher-order association networks (default mode, control), while structural differences concentrated in lower-order cerebellar and subcortical pathways. These findings provide a lifespan-wide, multimodal map of sex differences in brain networks which may help inform sex-specific vulnerability and resilience to brain disorders.

---

Sex differences are increasingly recognized as a critical factor in brain health and disease. Many neurological and psychiatric disorders show striking sex disparities in their prevalence throughout life. For example, males are about four times more likely than females to be diagnosed with autism spectrum disorder^1^, two to three times more likely to develop substance use disorders ^2^, and are more frequently affected by Parkinson’s disease^3^. Females show two times higher prevalence of anxiety^4^, 1.7-2 times higher prevalence of depression^5^, and a substantially higher prevalence of dementia in older age compared to age-matched men^6^. Moreover, certain conditions are exclusive to one sex and have higher risks during particular life stages; for example, postpartum depression is the most common psychological condition following childbirth, with the risk of depressive episodes being twice as high as during other periods of a woman’s life ^7^. However, despite increasing recognition that sex differences play a critical role in neuroscience and psychiatry, as well as growing recognition of “sex as a biological variable” as a research priority^8^, only limited studies have incorporated sex using optimal designs or treated it as a discovery variable^9^. As a result, how sex shapes or is shaped by the brain’s anatomical and physiological connections remains overlooked^9^. It is imperative to consider sex when investigating brain-behavior relationships as there may be differences in underlying disease mechanisms that in turn require different preventative approaches or treatments for optimal clinical care for all sexes^10^.

There is a body of work demonstrating robust sex differences in the human brain in terms of structure and function^11–13^. It should be emphasized that while males and females exhibit patterns of brain connectivity that are distinct when comparing across groups^14–16^, their distributions still possess significant overlap^17–19^. Sex differences in brain and behavior are not static; previous work has shown varied effects over the key developmental stages of puberty, reproductive maturity, and aging^13;18;20^. For example, functional connectivity (FC) in somatomotor, visual, control, and limbic networks were found to have the largest sex differences among children^20^, the default mode network was found to have largest sex differences in young adulthood^13;21^, and somatomotor and prefrontal/posterior cingulate were found to have the largest sex differences in an older cohort^22^. Recent work^23^ analyzed FC from human datasets across the lifespan and found that females had lower FC overall but also had higher variance, with female sensorimotor and limbic networks having lower segregation and the default-mode and ventral attention networks having larger segregation. However, this work only compared the overall lifespan sex differences by examining the entire sample with sex as a covariate, rather than investigating if sex differences change dynamically across the lifespan.

No work to date has performed a comprehensive comparison of male and female brain connectomes across the lifespan - from childhood to old age - to uncover the shifting relationship between sex and brain function as well as structure. A large-scale neuroimaging review^24^ summarized evidence for sex differences in brain structure over the lifespan, while emphasizing variability and methodological challenges in their interpretation. One study in a large cohort of children revealed larger brain volumes and higher proportions of white matter in males, and larger proportions of gray matter in females and lower mean/transverse diffusivity in primarily the superior corticostriatal white matter bundle in females^25^. Another study of individuals aged 8-22 showed very few sex differences in children, but prominent sex differences emerging in adolescence and continuing to young adulthood, namely that females had stronger across-hemisphere cortical structural connections and males had stronger within hemisphere cortical connections, an effect which was reversed in the cerebellum^26^. Another study^27^ analyzed T1-weighted anatomical scans across the lifespan and showed that males exhibit greater variance in brain structural metrics (subcortical volumes, cortical surface area, and cortical thickness) than females, with these differences in variance becoming less prominent with increasing age. These findings suggest the presence of dynamically changing sex differences across the lifespan, but, critically, stop short of a comprehensive analysis of how sex differences in brain networks change from childhood to old age.

There are many methodological issues to consider when associating brain network measures with some outcome, either demographics or behavior (i.e., brain-behavior mapping). First, there is currently no consensus on an optimal processing pipeline for fMRI and dMRI data when extracting connectomes, e.g., choices regarding atlas parcellation, artifact removal, and connectome metrics. To further complicate the issue, different connectivity ”flavors” resulting from these varied choices can lead to differing conclusions when performing brain-behavior mapping^28;29^. Second, connectome datasets are often high dimensional in terms of measured parameters^14^ and low in terms of observed samples from individuals (often N*<*100), a combination which can mean unreliable and non-reproducible results^30^. Our recently developed tool, Krakencoder, a multimodal connectome fusion and translation tool, can help mitigate these challenges ^31^. The Krakencoder generates a low dimensional “fusion” latent space that captures many structural and functional connectome flavors in a single compact and comprehensive representation, extracting common and useful information that can be robustly used for optimized brain-behavior mapping. Here, we further modified and retrained the Krakencoder to enhance the encoding of sex-specific information. Additionally, we leverage FC and SC data from a large set of 1286 participants aged 8-100 years compiled from the Human Connectome Project’s (HCP) Lifespan and Human Disease datasets in order to boost reliability of our findings^32–37^.

This paper provides a dynamic perspective on sex-based connectivity differences across the human lifespan by using both raw connectome data and the Krakencoder’s fused connectome representations from HCP studies to build statistical models classifying an individual’s sex. We leverage model interpretability to identify which brain networks and connections are most important to sex classification across the lifespan. Our results mapping trajectories of sex differences in FC and SC reveal how brain function and structure diverge in males and females from childhood to old age. These insights could help us understand sex-biased incidence and timing of neuropsychiatric and neurodegenerative disorders, and can also inform age- and sex-related diagnostics, therapeutics, and interventions.

## 1 Results

### 1.1 Sex classification accuracy varies across the lifespan

To study how sex-related patterns evolve with age, we divided individuals into ten age bins with approximately equal numbers of samples. Within each bin, three sex-classification models were constructed: (1) logistic ridge regression applied to the latent space of the original Krakencoder, (2) logistic ridge regression applied to the latent space of the sex fine-tuned Krakencoder, and (3) an ensemble logistic regression model trained directly on raw connectome data. Each approach was applied in three data settings, using: all 15 connectome flavors (Fusion), 9 FCs only (Fusion FC), or 6 SCs only (Fusion SC). The *Sex Fine-tuned Krakencoder* is the original Krakencoder that has been fine-tuned using data from 448 subjects (denoted the ”fine-tuning dataset”), but with an additional sex prediction decoder arm in the loss function. The remaining 838 subjects constituted the sex-classification dataset used for sex prediction (see Section 4.2.1 for details). Figure 1 (a-c) illustrates sex classification accuracy (i.e., % of the test set correctly classified) over the lifespan for our three models (Ensemble, Krakencoder, and Sex Fine-tuned Krakencoder), with each evaluated over the same 100 random train-test splits per age bin. Overall, the sex classification accuracy for models using both connectome types was quite high, ranging from 0.8-0.95 (*Sex fine-tuned Krakencoder*), with the single connectome modality models dropping modestly and fusion SC outperforming fusion FC at older ages. The original Krakencoder models do at least as well and many times significantly better than the ensemble model based on the raw connectome data (corrected p*<*0.01). The *Sex Fine-tuned Krakencoder* has significantly better accuracy than the ensemble model in every age bin and connectome modality (all corrected p*<*0.01), which robustly demonstrates the applicability of the Krakencoder’s latent space to be shaped such that it better represents some external variable.

**Figure 1:**
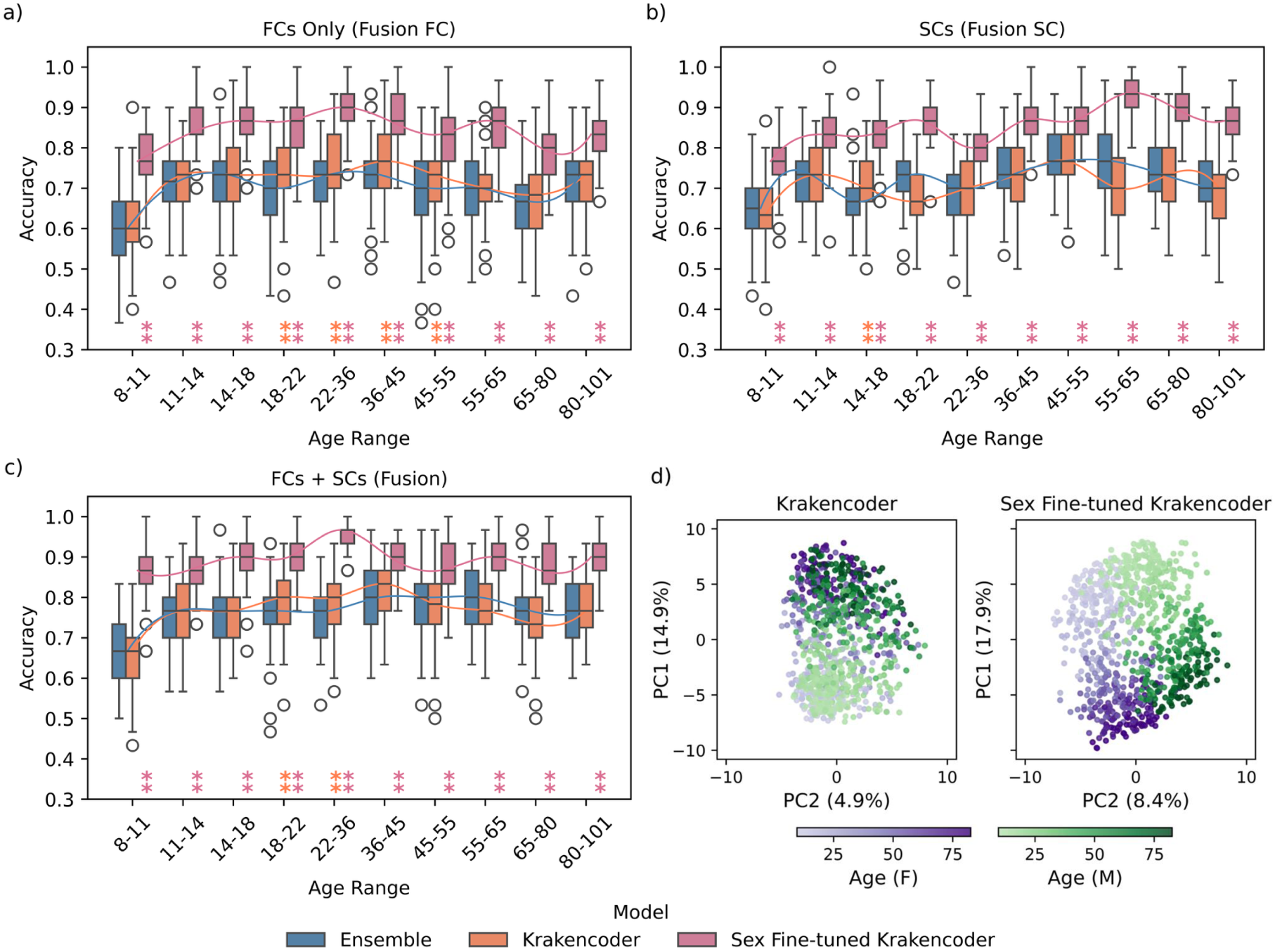
Sex classification accuracies across the lifespan using three different models (Ensemble in blue, Krakencoder in orange, Sex Fine-tuned Krakencoder in pink) with three different connectome modalities (a) Fusion FC, (b) Fusion SC, and (c) Fusion (FC and SC). For each model, a spline was fit to the average accuracy to visualize age-related trends in sex classification accuracy. One-sided permutation tests were conducted to evaluate whether the mean accuracy of each Krakencoder model is significantly higher than that of the ensemble model. P-values were corrected across the 10 age bins within each modality by FDR control (∗∗ corrected *p <* 0.01), where color indicates the model that had significantly better accuracy than the ensemble model. d) First two PCA projections (PC1 and PC2) of the low-dimensional latent space of the Krakencoder (left) and Sex Fine-tuned Krakencoder (right). Axes are labeled with the percentage of total variance explained by each component (e.g., PC1, xx%). Points represent individuals where color indicates sex (purple = female, green = male) and color intensity indicates age (lighter = younger).

All model types and connectome modalities demonstrate that sex prediction accuracy is lowest in early life (age 8-11), which indicates males and females exhibit the most similar FC and SC during the prepuberty or early pubertal period, with sex differentiation emerging during adolescence and persisting into adulthood. Models that use both SC and FC, as well as those based on FC alone contain peaks in young-to-middle adulthood (ages 22–45), with the sex fine-tuned Krakencoder reaching its highest accuracy at ages 22–36, followed by a gradual decline in later life. Models based only on SC achieve higher accuracy in middle-to-late adulthood (ages 36–80), suggesting ongoing sex-specific SC changes in middle to older ages. Although younger and older participants exhibited greater in-scanner head motion, quantified by mean framewise displacement (FD), than midlife (Supplementary Fig. S4), the two oldest age bins showed FD comparable to or exceeding that of the youngest while retaining relatively strong sex classification accuracy. Males and females did not have significantly different motion within any age bin, indicating it is likely not the primary factor driving accuracy fluctuations. Analyses of site effects (Supplementary Fig. S5) further showed minimal impact of data collection site effects on our results. Supplementary Fig. S2 showed that different classifiers predicting sex from the sex-fine-tuned Krakencoder latent space yielded similar lifespan patterns. We observed somewhat lower classification accuracy for the ensemble model compared with existing sex classification studies that rely on connectome data. We hypothesized that this reduction was driven by smaller sample sizes resulting from dividing the dataset into 10 age bins; therefore, we created a sex classification model using the full, unbinned classification dataset (Supplementary Fig. S3). All three models achieved high sex-classification accuracy, demonstrating that our sex fine-tuned Krakencoder is more robust to reduced training sample sizes than the ensemble model.

We applied PCA to the low-dimensional representations of Krakencoder and Sex Fine-tuned Krakencoder to visualize dominant sources of variation. Figure 1 (d) shows the projections of the first two principal components (PC1 and PC2) for subjects in the sex-classification dataset. In the Krakencoder, age varies primarily along PC1, while females and males are not clearly separable, at least in the first two PCs. In contrast, the Sex Fine-tuned Krakencoder exhibits a more structured latent space, with age-related trajectories still being captured along PC1 (despite the fact that age was not explicitly included in the training) and a clear separation between females and males along PC2. These results provide a visualization of the power of the Krakencoder to reorganize its low-dimensional representation to better represent external variables like sex. Sex specific lifespan trajectories in the latent space (PC2 representing sex versus age) in Supplementary Fig. S1 show patterns consistent with the main classification analyses, with the male and female lines of best fit being closest at the younger ages and farthest apart in midlife.

### 1.2 Importance of brain networks in sex classification

#### 1.2.1 Brain network importance in sex classification varies across the lifespan

We investigated brain network importance using a *Network Inclusion* sensitivity analysis, focusing on the 7 functional Yeo networks^38^, including visual (VIS), somatomotor (SMN), dorsal attention (DAN), ventral attention (VAN), limbic (LIM), control (CON), default mode (DMN), along with Subcortex (SUB) and Cerebellum (CBL). For a given target network or network pair, all other edges were replaced with that age bin’s training population mean (so-called masked data), then this masked data was used to predict sex. Sex classification accuracy on the model with the masked data was compared to accuracy of the model trained on the original (un-masked) data. Figure 2 presents the network-level and network-pair results from the *Network Inclusion* sensitivity analysis performed with the *sex-fine-tuned Krakencoder*. Across all three scenarios (Fusion, Fusion FC, and Fusion SC), a smaller accuracy drop suggests that the retained network, or network pair, contains more information used to classify sex.

**Figure 2:**
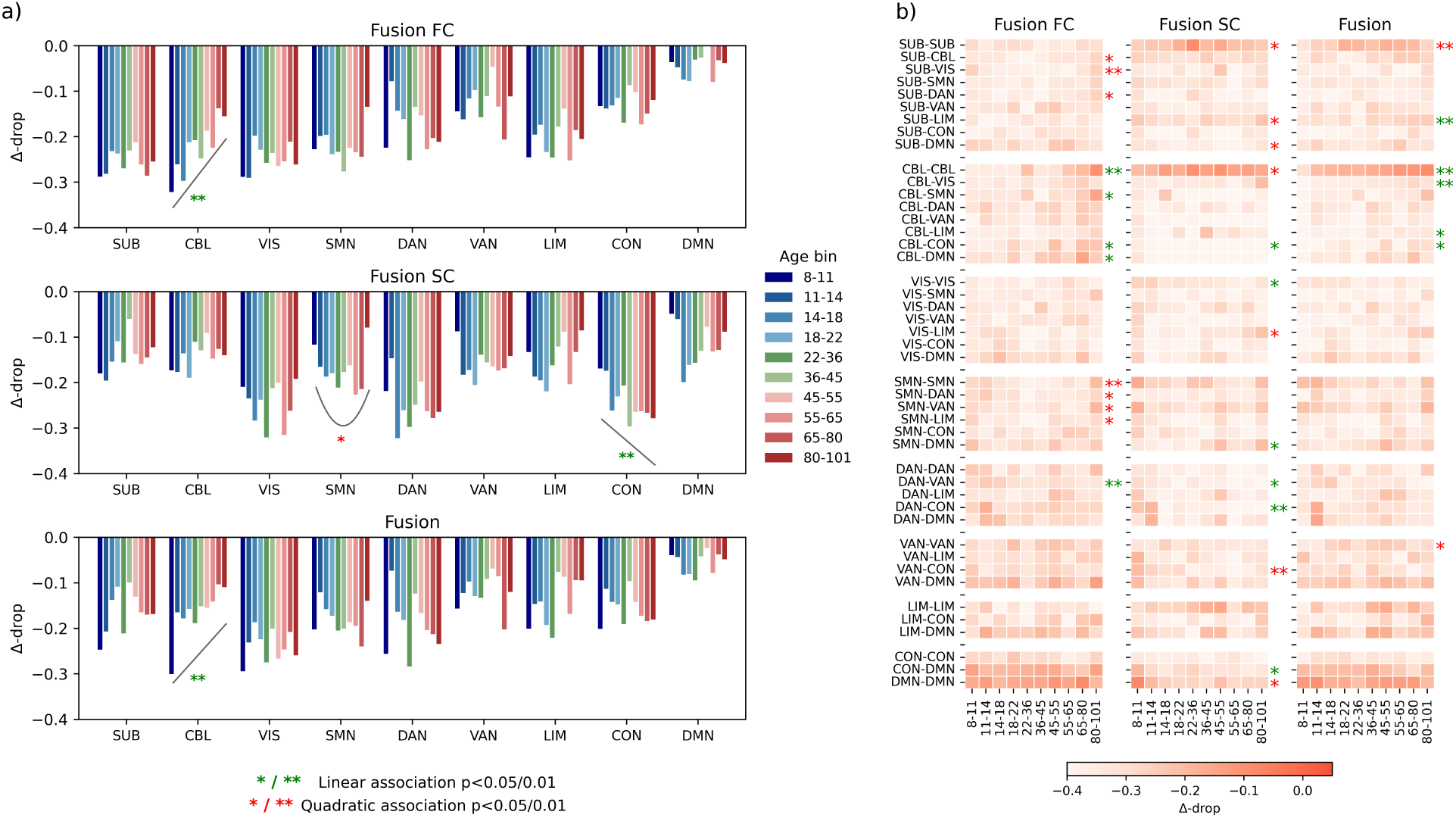
Sex classification feature importance. Average relative sex classification accuracy drop (Δ-drop) when only that network is included in the model; values closer to 0 indicate that the network is more informative of sex while more negative values indicate that the network is not as important for classifying sex. (a) Network-level analysis (y-axis: Δ-drop, x-axis: 9 networks). Curves show the trajectory of Δ-drop across age for each network for those with a significant monotonic association or a significant quadratic association across age bins. (b) Heatmaps showing network-pair contributions to sex classification, with darker colors indicating greater contribution of connections between regions in those two networks. Rows denote network pairs and columns denote age bins for the three sex-fine tuned Krakencoder models (fusion, fusion FC and fusion SC). Significant monotonic trends (Spearman’s rho test) across age are marked by green stars: ∗∗ for (uncorrected) *p <* 0.01 and ∗ for *p <* 0.05. Significant quadratic associations (a quadratic regression fit) with age are marked by red stars: ∗∗ for (uncorrected) *p <* 0.01 and ∗ for *p <* 0.05. These p-values are uncorrected, as they are used only for interpreting sex-classification feature importance changes over time. Figures S6 and S7 in Supplementary Materials provide the exact values of Δ-drops.

Figure 2 (a) presents the network-level contributions to sex classification accuracy. In FC (Fusion FC), higher-order networks, particularly DMN across all ages, and to a lesser extent CON and VAN, exhibited the smallest accuracy drops, indicating their strong and stable sex associations, with peak divergence in midlife (ages 45–55), a range which is centered around the typical timing of menopause in females (51-52). As people age, FC in the CBL becomes increasingly important for classifying sex in a linear manner. In contrast to FC, SC in lower order networks, namely SUB and CBL, along with DMN (as in FC), and to a lesser extent VAN and LIM, show highest importance in sex classification. SC in CON had significantly linearly decreasing importance for sex classification as age increases, while SMN had a u-shaped importance where it was most important for sex classification at young and older age and less important in mid-life. Fusion results are largely a combination of FC and SC features, with higher-order networks such as DMN, CON, LIM, VAN, and, to a lesser extent, SUB, CBL being the sex-informative. Among them, DMN is clearly most sex-informative. The CBL exhibits a linear decreasing association, indicating that functional and structural connections in this network may increase in their sex differences as people age.

Figure 2 (b) shows heatmaps of the relative accuracy drops for only including specific network-pair edges or connections. FC connections within and between higher-order networks (bottom rows), particularly between regions within the DMN (DMN-DMN), between regions in DMN with 1) control (CON-DMN), 2) limbic (LIM-DMN), and 3) ventral attention (VAN–DMN), exhibit the strongest influence on sex prediction accuracy across almost the entire lifespan. Connections within somatomotor (SMN-SMN) and between somatomotor and attention/limbic networks (SMN-VAN, SMN-DAN, SMN-LIM), exhibit significant quadratic trends, indicating stronger sex differentiation at younger and older ages. Connections with CBL, particularly within CBL, show significant linearly increasing feature importance for classifying sex with increasing age. SC between cerebellar regions and between subcortical regions were most important for sex classification, and to a lesser extent SC within limbic regions (LIM-LIM) and somatomotor regions (SMN-SMN). SC between regions of subcortex and between regions of the cerebellum had quadratic trends indicating that the sex differentiation for these networks was strongest in midlife. The fusion model’s network pair feature importances are largely a combination of the two other models, where connections within subcortical regions, within cerebellar regions, and within default mode network regions had the most sex differentiation. Connections within the cerebellum increased in sex importance linearly with age, while subcortical regions had largest importances in midlife compared to young and old ages. Supplementary Figures S6 and S7 provide detailed values of the relative accuracy drops underlying sensitivity analyses in Figure 2.

#### 1.2.2 Directionality of sex differences in network connections

Next, we investigated the feature importances of the logistic regression models built directly on the raw connectome data in order to 1) replicate our feature importance findings from the Krakencoder and 2) shed light on the directionality of the sex differences that the Krakencoder feature importances do not explicitly provide. Figures 3 and 4 show the average standardized Haufe-transformed coefficients across the 9 FC models and 6 SC models, respectively, for the 9 networks of interest split over left (LH) and right hemispheres (RH). Haufe-transformed coefficients^39^ more directly reflect underlying sex-related signals in the presence of strong collinearity. Each circle plot represents the feature importances for a given age bin, where the Haufe transformed standardized coefficients are first averaged across the 100 replications of the models per flavor, then averaged across SC or FC flavors (see Section 4.2.2). Panel (a) shows the connections where females had stronger connectivity (darker blue = larger in females vs males) and panel (b) shows the connections where males had stronger connectivity (darker orange = larger in males vs females). Link thickness encodes the magnitude of network-pair importance, offering an intuitive visualization of the directionality and magnitude of sex differences in FC or SC across brain networks, hemispheres, and age groups. To emphasize robust effects, only the top 5% strongest network-pair connections per age group are shown. Results using a relaxed 15% threshold are provided in Supplementary Figure S9, allowing a deeper dive into feature importances under less restrictive thresholding.

**Figure 3:**
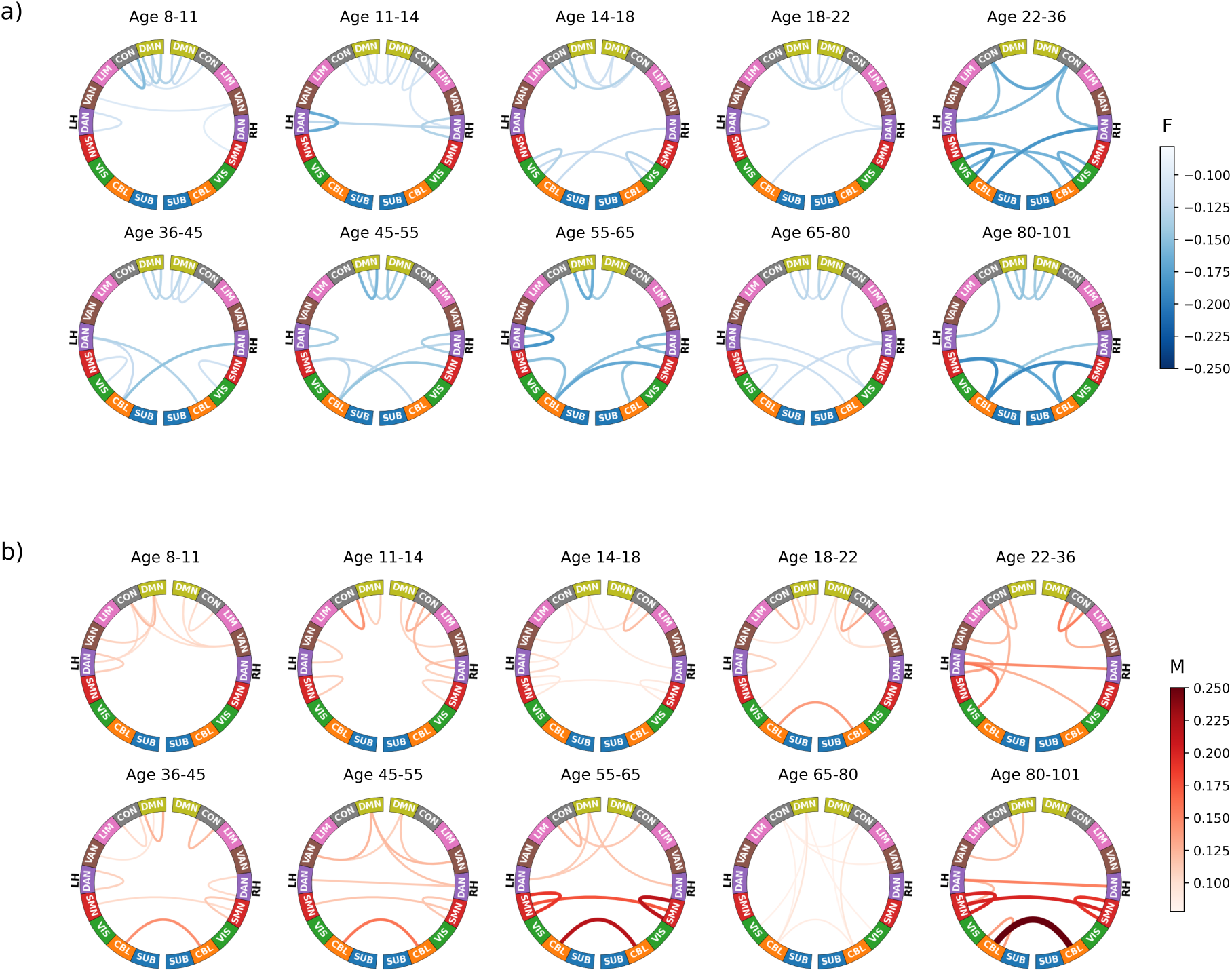
Sex differences in FC across the lifespan, where each circle represents the feature importance of the sex classification model from a specific age bin. (a) Network connectivity feature importances where larger FC was associated with female sex. (b) Network connectivity feature importances where larger FC was associated with male sex. Darker colors / thicker lines indicate greater feature importance when predicting sex. LH: left hemisphere, RH: right hemisphere, F: female, M: male. In order to emphasize the most salient effects, all chord diagrams display only the top 5% strongest connections within each age group, based on the magnitude of the average standardized Haufe coefficients in the matrix of networkpair averages.

**Figure 4:**
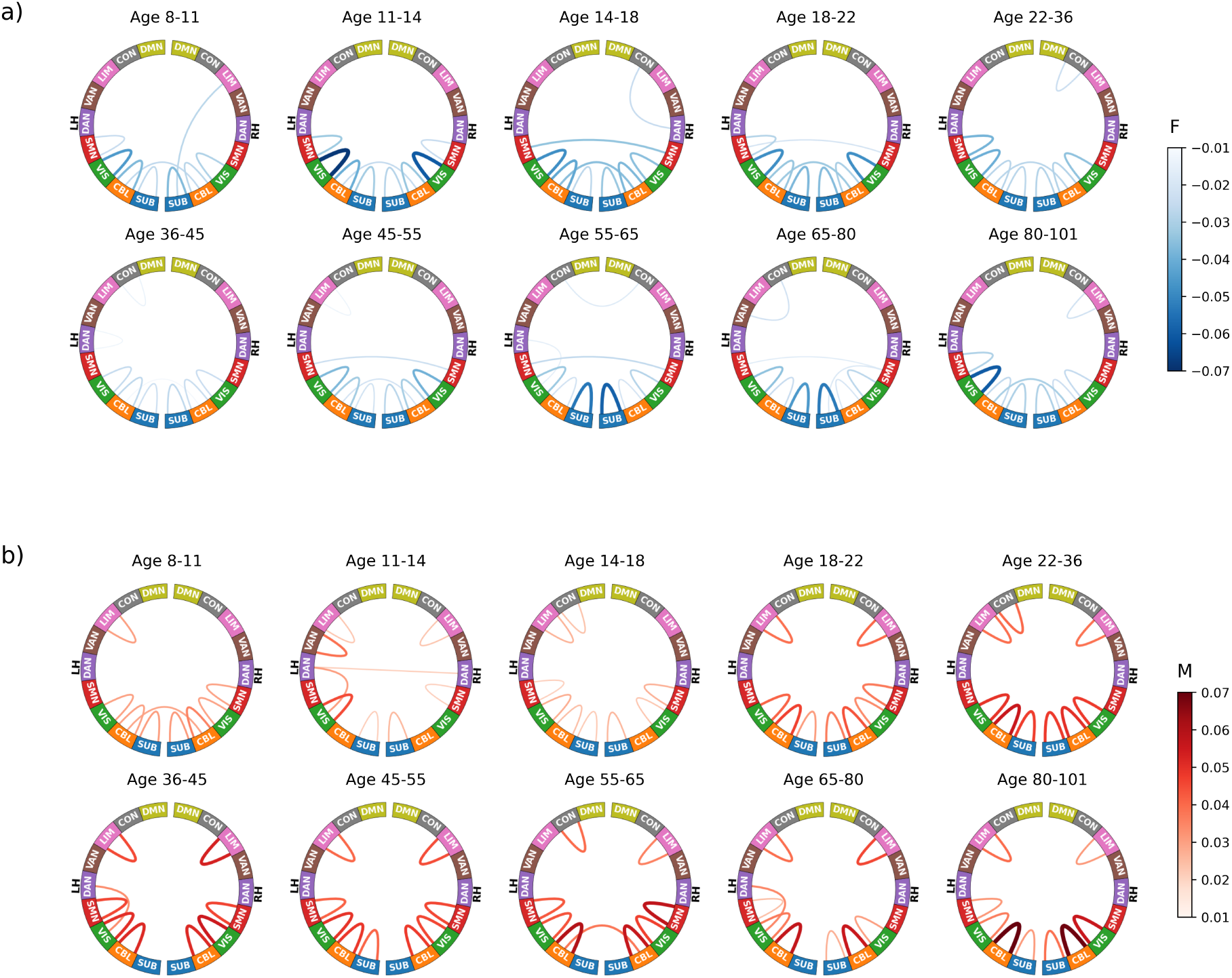
Sex-specific feature importance patterns for SC across the lifespan, where each circle is an age bin. Panel (a): Network connectivity feature importances where larger SC was associated with female sex. Panel (b): Network connectivity feature importances where larger SC was associated with male sex. Darker colors / thicker lines indicate greater importance. LH: left hemisphere, RH: right hemisphere, F: female, M: male. Only the top 5% strongest network-to-network connections are shown for each age.

Overall, FC in higher order networks are different between the sexes consistently across all ages, while FCs in lower order regions, particularly within sensorimotor networks and cross-hemisphere cerebellar regions, are increasingly sex differentiated as people age. Generally speaking, female sex is associated with stronger FC between regions of the DMN (DMN-DMN) across the lifespan. Stronger FC between left CBL and right DAN appears to emerge in females around 14, and this pattern persists throughout the rest of the lifespan. Left CBL to bilateral SMN FCs are stronger in females starting around midlife and continuing through old age, as further confirmed via a significant linear increase in this network-pair feature importance with age (see Supplemental Figure S10). Generally across the lifespan, males had stronger FC within CON regions (CON-CON) and from CON and DMN to attention networks (DAN/VAN). While females had stronger FC between CBL and SMN/DAN, males had stronger across-hemisphere CBL and across-hemisphere SMN connections starting around late adolescence, continuing to become more sex differentiated with age as validated via a significant linear trend in Figure S10. Importantly, these results generally align with the Krakencoder’s feature importances in Figure 2, demonstrating the stability of our conclusions. Together, these observations show that FC within and between higher order networks (DMN, CON and attention networks) are most sex differentiated regardless of age range, and, that lower order (CBL and SOM) FC becomes increasingly sex differentiated as people age.

Figure 4 visualizes the directionality of the SC connections that drive sex classification across the lifespan. Feature importances are primarily concentrated to within-hemisphere and within-network SCs of lower order networks, i.e., cerebellar, visual, subcortical, and somatomotor (see also Figure S8). Interestingly, each sex exhibited different regions within these ipsilateral networks that had larger/smaller values compared to the other sex. A more detailed region-wise sex differences matrix is given in Supplemental Figure S11. One exception to these broader observations is that females had stronger SC in across-hemisphere SMN at various ages (panel a), while stronger SC within ipsilateral regions of the SMN was associated with male sex (panel b). We also observe that ipsilateral cerebellar SCs became increasingly stronger in males compared to females beginning in late adolescence (*>*18) and into older age, reflected by significant linear increases in male coefficients (and significant decreases in female coefficients) over the lifespan (Figure S10). Again, these feature importances echo those identified by the sex-fine tuned Krakencoder, which highlights our feature’s reproducibility.

## 2 Discussion

Sex is increasingly recognized as a key variable for understanding trajectories of brain development, brain aging, and risk for neurological and neuropsychiatric conditions. Here, we show that a basic organizational feature of the brain - the structural and functional connectomes - varies across the lifespan in a sex-dependent manner. Leveraging our new AI-based Krakencoder, we found that sex differences in the brain’s structural and functional connectome are minimal in early childhood, pronounced in young to mid-adulthood, and diverge by modality later in life. Higher-order functional connections (default mode, control, and attention networks) and lower-order ipsilateral, within-network structural connections (cerebellum, visual, subcortical and somatomotor) drove sex classification. While functional connectivity in higher-order networks was sex-divergent across the entire lifespan, other networks, such as those within the cerebellum, became increasingly sex-divergent over time. Specifically, females tended to have stronger functional connections between default mode regions regardless of age, while males exhibited stronger FC between regions of the default mode and attention networks and within ipsilateral regions of the control network. Males had increasingly stronger FC compared to females between bilateral cerebellums as they aged, while females had increasingly stronger FC between cerebellum and somatomotor networks as they aged. In terms of the brain’s structural organization, ipsilateral, within-cerebellar SCs grew stronger in males compared to females as they aged. To our knowledge, these results represent the first analysis detailing how sex shapes the brain’s structural and functional connectome across the lifespan, advancing ongoing efforts to integrate sex difference analyses into brain research^9;40^.

Our sex classification accuracy trajectories mirror overall sex hormone level trajectories across the lifespan, suggesting that sex hormones likely drive some of the observed effects ^41^. Individuals older than six months exhibit low and largely indistinguishable sex hormone levels until puberty, likely contributing to our finding of the lowest levels of sex classification accuracy during early childhood. Despite the relatively lower accuracy, it remained better than random chance, indicating that sex differences in brain structure and function are present even before the pubertal increase in sex hormones production^20;42;43^. These early sex differences could be genetic, social, or environmental in origin, a possibility that exceeds the scope of the current study. During the reproductive years, between puberty and midlife, sex hormones reach their peak in both sexes, coinciding with our highest observed sex classification accuracies. From mid to late life, sex hormones drop sharply in females due to menopause and gradually in males with the onset of andropause, corresponding with a reduction of sex differences in FC, which may be more impacted by hormone shifts than SC. In contrast, sex classification accuracies of SC continue to increase from middle to later life, potentially reflecting an accumulation effect of continued social, environmental and/or biological/hormonal exposures influencing white matter organization divergence between sexes. Higher order network FCs had the largest sex differences across the lifespan while the cerebellum had increasingly sex divergence with age. Most prominently, females had stronger FC within regions of the DMN across the lifespan, while males had stronger FC between DMN-attention networks and within ipsilateral regions of the CON, reflecting previous work^18;44;45^. Alterations in default mode connectivity have been associated with neuropsychiatric disorders like depression and anxiety (more female-prevalent)^46–49^. The perimenopausal transition in females as well as aging-related endocrine or metabolic changes in both sexes could modulate sex differences in brain structure, connectivity, and networks^50–52^. One study^53^ showed that sex differences in certain networks (namely the DMN) become prominent around midlife. In our feature importance, the DMN is maximally important in the 45-55 age range, centered around the average age of menopause (51-52). Menopause has also be linked to pronounced cerebellar FC differences, including lower FC within CBL, and stronger FC between CBL-frontal and CBL-striatum in late postmenopausal females (aged *>* 54)^54^. This pattern aligns with our sex differences in CBL FC that emerge in middle adulthood and increase with age, wherein females had stronger FC between CBL and cortical networks (DAN and SMN) and males had increasingly stronger FC interhemispheric CBL connectivity from midlife to older age. These findings suggest that cerebellar reconfiguration occurs around the menopausal transition, consistent with previous findings of volume changes in this structure that vary with hormone shifts in menopause ^55^. The convergence of our connectivity findings with the timing of perimenopause in women and gradual androgen changes in men supports the interpretation that hormonal and developmental processes play a role in these midlife shifts in sex differences. Prior studies have shown that hormones, sex, midlife aging, and menopause are associated with memory^56^, cognition, or dementia^57–59^. Indeed, endocrine aging in women (particularly changes in estradiol levels) has also been linked to reorganization of brain circuits and a higher susceptibility to female-prevalent disorders like depression and Alzheimer’s disease^60^. Such midlife divergences may reflect the impact of changing sex hormone levels on brain circuitry or accumulated life experience shaping brain structure.

SC in lower order networks, including the subcortex, cerebellum, visual, and somatomotor networks had the largest sex differences, consistent with several prior works. For example, males aged 8-22 exhibited stronger SC than females between the left cerebellar hemisphere and the right cerebral cortex^26^ . Among individuals aged 22–35 years, diffusion-derived microstructural parameters exhibited the greatest sex differences within motor and limbic systems^61^. Our work extends prior findings across the lifespan and suggests that sex-associated neurodevelopmental and hormonal trajectories may contribute to age-dependent differences in SC in lower-order ipsilateral networks, with distinct timing and patterns observed between females and males. Furthermore, our findings of males having generally stronger SC between regions of the same network and hemisphere, with females having stronger across-network and -hemisphere SC aligns with previous work showing higher modularity of male vs female SC networks^26^. Finally, we found generally more consistent sex-specific features across the lifespan (compared to FC sex differences), which appeared to become even more sex divergent across aging. This indicates perhaps a weaker relationship between the structural connectome and lifespan changes in hormones or environmental factors compared to the functional connectome, which is aligned with the more static nature of SC generally.

Here we analyzed sex differences within discrete age bins, allowing for fluctuations in sex differences across the lifespan. Most previous connectome studies model age and/or sex as covariates, which limits the ability to capture age-dependent changes in sex differences^23;62;63^. While models that treat age as a continuous variable and include sex×age interaction terms^53;62^ are statistically powerful for testing gradual, sex-specific age effects, they impose smooth (e.g., linear or quadratic) assumptions on how sex differences evolve across the lifespan. Such parametric formulations risk obscuring stage-specific transitions that occur in puberty, adolescence, reproductive years, menopause, and aging, all of which are known to impact brain network organization^42;52;64^. For example, both^53;62^ test the significance of the interaction between age and sex to explore sex differences in age trajectories, which only allow sex differences vary linearly (or quadratically) with age. In contrast, our approach avoids parametric age assumptions, allowing sex-related effects to vary across the lifespan without imposing assumptions on their functional dependence on age. Many existing connectome studies investigating group differences (like sex differences) rely on traditional statistical comparisons, such as group mean and variance differences^18;27^, and at times employ traditional statistical and machine learning prediction models^14;65;66^, which are limited in capturing complex, non-linear connectivity patterns. In contrast, more advanced machine learning techniques^13;67;68^ can uncover latent patterns of sex differences overlooked by classical methods. Using our advanced AI-based tool (the Krakencoder), we demonstrate that there are indeed robust, distributed multivariate connectome patterns that can distinguish the sexes across the lifespan. We demonstrate the advantages and flexibility of our Krakencoder tool in fusing different flavors of connectome processing, and different modalities, in a way that increases SNR, reduces connectome dimensionality, and maximally explains some external variable simultaneously.

While our study provides novel insights, several potential issues and limitations must be addressed. First, in-scanner motion might influence sex classification accuracy differently across the lifespan. Although younger and older participants generally exhibited greater motion compared to midlife participants, sex classification remained accurate even in high-motion age bins and did not differ between males and females within age groups, suggesting minimal impact of motion on the results. Second, the MRI data was collected across several different locations which can cause bias in extracted connectome values ^69;70^. Our analysis that adjusted connectome data for potential site effects (see *Site Effects* in Supplementary Figure S5) demonstrates that the impact of site differences on our sex classification findings are also minimal. Additional limitations include the cross-sectional rather than longitudinal design and the lack of connectome data for children younger than 8 years old due to MRI quality issues (excessive motion). Recent studies report that sex differences in the brain can emerge even before birth^71^ or at birth^72^, so our future work will exploit public datasets from this age range as well. Finally, and perhaps most importantly, we did not investigate the impact of gender identity, pubertal stage, past pregnancies, menstrual cycle stage, menopausal status, oral contraceptive use, hormone levels, or other nuanced biological or environmental factors that could refine the understanding of sex differences. We chose to start here at the coarsest view - sex differences across the lifespan - which remains incompletely understood. However, it is clear that reproductive health and events (e.g., pregnancy or menopause) can impact brain connectomes. For example, it has been shown that women’s reproductive history and cumulative hormone exposure have measurable impacts on brain aging – multiple childbirths are associated with less apparent brain aging, whereas higher lifetime estrogen exposure is linked to more evident brain aging^73^. Additionally, fluctuations in functional and structural networks have been shown across the menstrual cycle^74–77^ and large shifts in brain volume have been reported across pregnancy^78;79^. Prior work also suggests that gender (different from biological sex used here) also impacts brain development in children^20^ and adults^80^. We pause here to highlight that, in most cases, reproductive health history information simply does not exist in neuroimaging datasets. We therefore emphasize the need for collecting full reproductive health history in a standardized way when carrying out neuroimaging studies across any age cohort, and are ourselves taking strides to create such a standard^81;82^.

This study provides the first analysis of how sex differences in brain networks evolve across the human lifespan. Functional and structural connectivity networks exhibit complementary, age-dependent trajectories: functional differences are dynamic, peak in midlife, and primarily involve higher-order networks, whereas structural differences are are high in midlife and continue to diverge with aging, particularly within lower-order networks. These findings underscore distinct sex-specific organizational principles across functional and structural domains and highlight the importance of incorporating reproductive health measures to better understand brain–behavior relationships and sex-specific disease mechanisms. Understanding how sex shapes brain connectomes across the lifespan is essential for elucidating brain–behavior relationships and sex-specific mechanisms of brain disorders, which in turn can advance personalized clinical care.

## 3 Acknowledgments

This work was supported by the Ann S. Bowers Foundation, through the Ann S. Bowers Women’s Brain Health Initiative (A.K., K.H.), and an NIH grant R01MH137166 (A.K.). Data were provided by the Human Connectome Project (HCP), WU-Minn Consortium (Principal Investigators: D. Van Essen and K. Ugurbil; 1U54MH091657) funded by the 16 NIH Institutes and Centers that support the NIH Blueprint for Neuroscience Research; and by the McDonnell Center for Systems Neuroscience at Washington University. Research reported in this publication was supported by the National Institute Of Mental Health of the National Institutes of Health under Award Number U01MH109589, and the National Institute On Aging of the National Institutes of Health under Award Number U01AG052564. The HCP-Development 2.0 Release data used in this report came from DOI: 10.15154/1520708. The HCP-Aging 2.0 Release data used in this report came from DOI: 10.15154/1520707. The authors thank F.-C. Yeh (Department of Neurological Surgery, University of Pittsburgh, Pittsburgh, PA, USA) for sharing preprocessed diffusion MRI data for HCP-Development and HCP-Aging subjects.

## 4 Methods

### 4.1 Data description and processing

The dataset we used comprises MRI, demographic, and behavioral data from 1286 healthy controls (643 female, aged 8-100) compiled from the HCP Lifespan studies — HCP-Development^32^ (HCP-D, ages 8–21) and HCP-Aging^33^ (HCP-A, ages 36–100+) — complemented by data from young adults (ages 22–35) who were healthy controls in three HCP disease studies ^35–37^.

Specifically, we include 543 children or adolescents from HCP-D (255 female, aged 14.76±4.12 years), 83 young adults (40 female, aged 26.34 ± 3.27) from three HCP disease studies (Early Psychosis^37^, Anxious Misery^35^, and Treatment-Resistant Depression^36^), and 660 older adults from HCP-A (348 female, 61.72 ± 16.04 years). Anatomical, resting-state fMRI and dMRI from all studies were acquired using the harmonized HCP-Lifespan protocol, and the images were preprocessed using identical pipelines and functional and structural connectomes extracted for 3 different atlases (fs86, shen268 and coco439) using different methods, as described previously^31^ and summarized in *Data processing pipeline* section of the Supplementary Materials. The end result is 15 connectomes for each individual, 9 functional and 6 structural. There are 3 types of functional connectomes (FCcorr - Pearson correlation FC without global signal regression, FCgsr - Pearson correlation FC with global signal regression, FCpcorr - partial correlation FC) for each of the three atlases and 2 types of structural connectomes (deterministic tractography - sdstream and probabilistic tractography - ifod2act) for each of the three atlases.

### 4.2 Sex classification models

We split this so-called “lifespan” dataset into 10 age bins with approximately the same number of individuals (see Figure 5d). Within each of the 10 age bins, the lifespan data was used to train and evaluate models that classify biological sex (male or female) based on brain connectomes (see detailed descriptions below in subsections 4.2.1 and 4.2.2) and Figure 5. The sex classifiers were created using three logistic ridge regression models trained on 1) ensembling the various flavors of raw connectome data, 2) the original Krakencoder’s latent space representation of the connectomes, 3) the fine-tuned version of the Krakencoder’s latent space designed to better represent sex differences within the lifespan dataset. To assess the importance of the different connectivity modalities in sex classification accuracy, each of the 3 models was trained on three versions of the data: all 15 connectivity flavors, 9 FC flavors only, and 6 SC flavors only.

**Figure 5:**
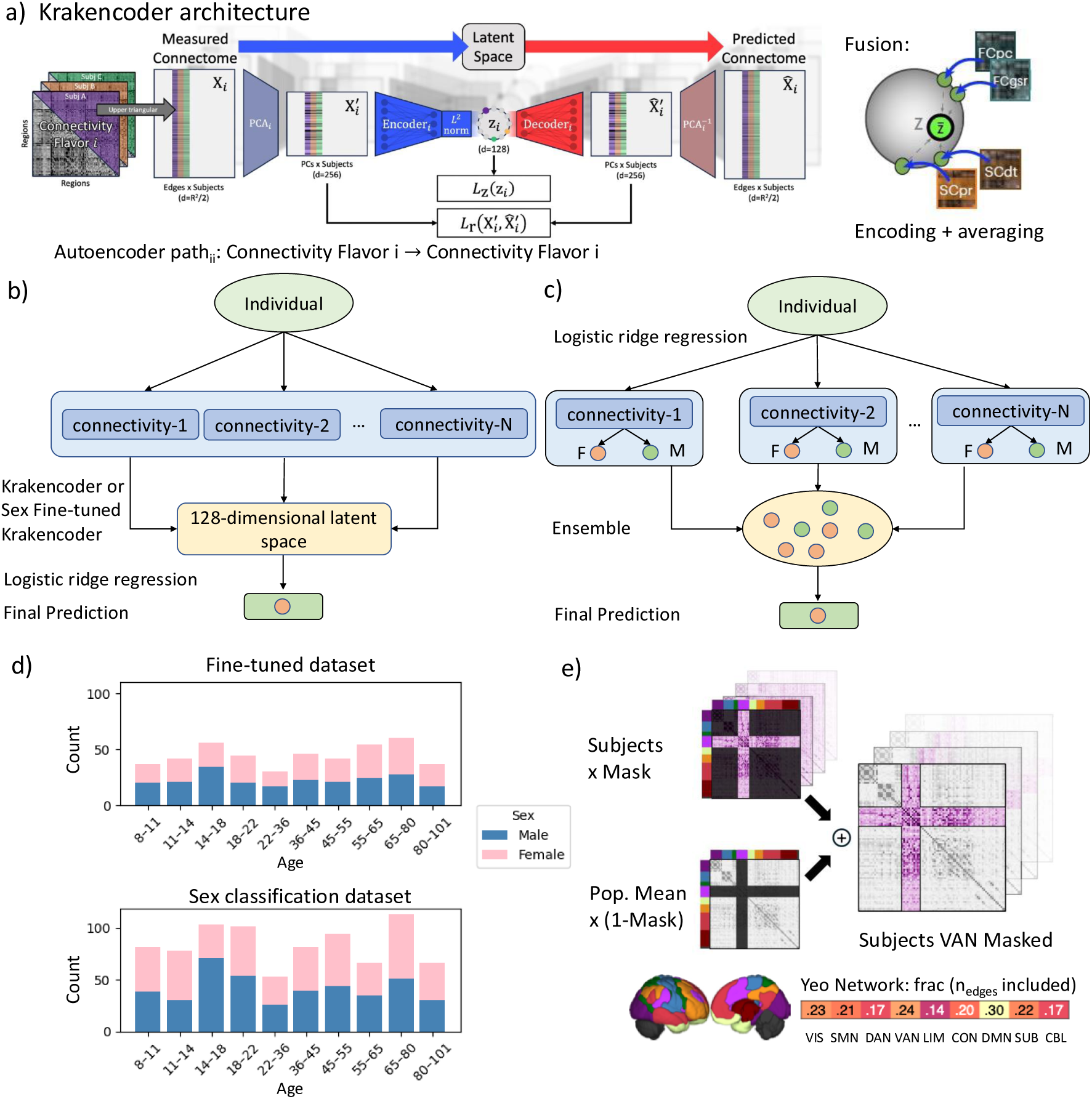
Overview of methods and datasets. (a) Krakencoder architecture: Each autoen-coder maps a connectivity type to a latent space (e.g., *z_i_* is the latent space for autoencoder mapping connectivity type i to itself). These latent spaces are integrated into a shared lowdimensional space *z̄*. *L_z_*and *L_r_* are the loss functions^31^. (b) The Krakencoder approach (original Krakencoder and Sex Fine-tuned Krakencoder): FCs and SCs (N=15); FCs only (N=9), or SCs only (N=6) are mapped into a 128-dimensional latent space and classified with logistic ridge regression. (c) Ensemble approach: separate regressions are trained per connectivity type and aggregated. (d) Sex distribution across age bins for the fine-tuning and sex classification datasets (fine-tuning set was used to fine-tune Krakencoder; sex classification set was used to predict sex). (e) *Network Inclusion* sensitivity analysis workflow (e.g., only retaining edges in VAN).

#### 4.2.1 Classifying sex with the Krakencoder’s latent space connectome representations

The Krakencoder is a multi-modal connectome fusion model^31^ that projects all connectivity flavors into a shared, 128-dimensional latent space. Each of the low-dimensional representations from the original connectome flavors can be averaged to obtain a “fusion” latent space representation of the FC (called fusion FC), SC (called fusion SC) or both (called fusion) (Figure 5 (a)). The Krakencoder was initially trained on the original HCP-Young Adult (HCP-YA, ages 22–36) data^83^, which differs in acquisition and age range from the lifespan dataset. To improve generalizability to the lifespan data and add specificity to sex classification, we fine-tuned the Krakencoder using 448 subjects (223 female) drawn from our lifespan dataset, with approximately equal sample size across age bins, see Fig 5 (d) top panel. This fine-tuning procedure also included an additional decoder, i.e., a single layer (128x1), to decode sex from the Krakencoder’s latent space. This decoder was added to the original Krakencoder’s connectome prediction paths (15 encoders, 15 decoders) to create a 240-path model (15 encoders, 16 decoders), where sex is predicted from the latent space of each of the 15 connectome inputs separately during training. The addition of this extra decoding arm resulted in a fine-tuning procedure that both ensured the Krakencoder’s applicability to the lifespan dataset as well as encouraged the latent space to contain information that could be used to classify an individuals’ sex.

All 1286 HCP-lifespan/disease subjects were used to build the sex classification models via logistic regression (see Figure 5 (b)), with the 448 subjects used for Krakencoder fine-tuning included only in the training splits for sex classification to avoid data leakage. The remaining 838 subjects were used for sex classification across all models. For each age bin, we randomly selected 30 subjects (15 female, 15 male) from the subjects’ data not used in fine-tuning the Krakencoder (so-called sex classification dataset), so that each bin had an equal number of test samples. For the training set, we combined the rest of the unused sex classification dataset samples within that bin with subjects from the 448 fine-tuning set of the same age range, and then downsampled to 30 training subjects (15 female, 15 male) in each bin. This process was repeated 100 times with different subgroups of training and testing subjects; test accuracy is reported for model assessment. This design ensured each age bin contained an equal number of subjects for both training and test sets, with a balanced number of females and males. The number of subjects was set to 30 for training and test as there were only about 50 subjects in the 22-36 age bin and we wanted to perform multiple cross-validation splits within each age bin, keeping the number of training samples consistent across age bins. Figure 5 (d) shows the distribution of fine-tuning and sex classification datasets.

Sensitivity analysis for the Krakencoder latent space models was conducted to identify which brain regions are most strongly associated with sex. Regions were grouped into seven cortical resting state networks according to Yeo^38^, including visual (VIS), somatomotor (SMN), dorsal attention (DAN), ventral attention (VAN), limbic (LIM), control (CON), default mode (DMN), along with subcortical (SUB) and cerebellar (CBL) networks. Within each age bin, we performed the previously developed *network inclusion* sensitivity analyses (Figure 5 (e)) using the *sex fine-tuned Krakencoder*. To perform this sensitivity analysis, every connectome edge was set to the population mean of the training set for that specific age bin, except for those involving regions within the network (or network-pair) of interest. The resulting masked data were used to predict sex (using *Sex fine-tuned Krakencoder*, followed by retraining a logistic regression model on the resulting low-dimensional representations), and the classification accuracy was compared to that obtained from the original (unmasked) data. A smaller relative drop in accuracy indicates that the target network (or network pair) retained more sex-relevant information, suggesting that this network (or network pair) is more informative for the sex classification. This enabled us to examine which inter- or intra-network connections are most informative for distinguishing sex and how these associations evolve across the lifespan. Because the networks differ in terms of the number of regions assigned to them, we wanted to make sure to report the % retained connections for each network’s inclusion analysis (see Figure 5e bottom). All networks are within the same range of retained connections, with most networks having around 20% retained. The limbic network had the smallest retained connections at 14% while the default mode network had the largest retained connections at 30%.

#### 4.2.2 Classifying sex with the original connectome data

Within each age bin, we built three ensemble models predicting sex from original (raw) connectome data: one combining all 15 connectome flavors, one with only the 9 FC flavors, and one with only the 6 SC flavors. For fair comparison, we used the exact same train and test cross-validation splits and number of repetitions as the models built on the Krakencoder’s latent space. Ensemble predictions were obtained via majority voting across the set of connectivity flavors being considered, resulting in 100 ensemble accuracies (one per random train-test split) per age bin. Figure 5 (b) illustrates the ensemble procedure across connectome flavors.

For interpretability, regression coefficients (*ω*_ridge_) were recorded and transformed using the Haufe transformation^39^. The Haufe transformed coefficients *ω*_haufe_ enhance interpretability by more directly reflecting the contribution of each feature, especially when collinearity among the input variables exists, as is the case with connectome data. Let *x* and *y* denote the input features and binary response, respectively. The Haufe transform is defined as *ω*_haufe_ = *c*Σ*_x_ω*_ridge_ ^39^, where Σ*_x_* is the input covariance matrix of *x* and *c* = var(*p*_logit_) with *p*_logit_ = logit(*P* (*y* = 1)) = *xω*_ridge_. To address shrinkage effects from varied regularization parameters that result in different magnitude regression coefficients across the models, we standardized the Haufe-coefficients as:

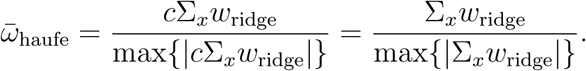

Suppose the input dimension is *d*, then the dimension of Σ*_x_* is *d* × *d*, and *ω̄*_haufe_, *ω*_haufe_, and *ω*_ridge_ are all *d* × 1. The operator max(|*v*|) gives the maximum absolute value of a vector *v*, so the dominator max{|Σ*_x_w*_ridge_|} is a scalar. Finally, to obtain reliable feature importance estimates, the standardized Haufe coefficients were averaged over 100 repetitions for each connectivity flavor.

