## Supplementary Information for "The progression of sex differences in brain networks across the lifespan"

### Data processing pipeline

The HCP-Development (HCP-D, aged 8-21) and HCP-Aging (HCP-A, aged 36-100+) data come from the HCP Lifespan studies[1–3], which differ from HCP-YA in both age range and acquisition parameters, with Lifespan data acquiring less than half as much fMRI and dMRI data as HCP-YA. For Lifespan HCP data, resting-state fMRI data were collected using 2.0 mm isotropic voxels, TR = 800 ms, and anterior–posterior (AP/AP) phase encoding, yielding a total acquisition time of 25.5 minutes. Diffusion MRI was acquired at 1.5 mm isotropic resolution with b-values of 1500 and 3000 s/mm$^2$, each sampled with 90 directions per shell, using anterior–posterior/posterior–anterior (AP/PA) phase encoding, for a total scan time of 22.5 minutes.

To complete our lifespan dataset, we needed data from individuals aged 22-35 that were acquired under the same protocol and MRI acquisition parameters as the HCP-D and HCP-A. Therefore, we included data from young adult healthy controls scanned as a part of three HCP disease studies, specifically the "Connectomes Related to Human Disease" (CRHD) studies that used the same protocol as HCP-D and HCP-A and were centrally coordinated and preprocessed by the "Connectome Coordinating Facility" (CCF). Three studies had fully preprocessed healthy controls in the young adult age range: Dimensional Connectomics of Anxious Misery (DCAM)[4] had 31 subjects (23 female, aged $26.52 \pm 3.82$), Early Psychosis (EP)[5] had 38 subjects (11 female, aged $26.05 \pm 2.19$), and Perturbation of the Treatment of Resistant Depression Connectome by Fast-Acting Therapies (PDC) had 14 subjects (6 female, aged $26.18 \pm 2.99$)[6]. We downloaded the fully preprocessed MRI data for these participants and applied consistent post-processing procedures as used for the HCP-Lifespan studies, including additional denoising, temporal filtering, and parcellation with fmriclean, and tractography reconstruction with MRtrix. This integration allows for a more continuous and age-complete analysis of structural and functional connectivity patterns across the human lifespan. The construction of functional and structural connectomes are exactly the same as that in[7], we provide details below.

#### Construction of functional connectomes

All resting-state functional MRI (fMRI) time series were preprocessed by HCP using the minimal preprocessing pipeline[8], which includes motion correction, susceptibility distortion correction, coregistration to anatomy and nonlinear template space (FMRIB Software Library (FSL) MNI152 template, 6th generation), as well as automated denoising using ICA-FIX[9]. We used a custom post-processing pipeline to identify motion and global signal outlier timepoints (global signal $> 5\sigma$, motion derivative $> 0.9$ mm), regress out tissue-specific nuisance time series (five eigenvectors each from eroded white matter and cerebrospinal fluid masks[10]) and motion-related time series (24 in total, including six motion parameters, their backward derivatives, and the squares of both[11]), and temporally high-pass filter ($> 0.01$ Hz for HCP-YA, $> 0.008$ Hz for HCP-Lifespan, using a discrete cosine transform projection). Outlier timepoints were excluded from nuisance regression and temporal filtering. Regional time series for each brain region for a given gray matter parcellation were obtained by averaging voxels in the denoised time series data. Regional time series for each of the four (HCP-YA) or two (HCP-Lifespan) scans were variance-normalized and concatenated, and Pearson correlation (FCcorr) between regional time series (excluding outlier timepoints) resulted in a region × region resting-state FC matrix for each subject. We also computed an FC matrix with global signal regression (gsr), by regressing the mean gray matter time series and its temporal derivative from each regional time series before computing region × region Pearson correlation (FCgsr). Finally, we computed a Tikhonov-regularized partial correlation (FCpcorr) that minimizes the average Euclidean norm between subject FCpcorr and the population mean of the unregularized precision matrices[12]:

$$\overline{\Omega} = \frac{1}{n_{\text{subj}}} \sum_{a=1}^{n_{\text{subj}}} FC_a^{-1}, \quad \text{1. Compute population mean of unregularized inverted FC}$$

$$\widehat{\Omega}_a = \left(FC_a + \lambda I\right)^{-1}, \quad \text{2. Find } \lambda \text{ that minimizes mean squared error (MSE) loss}$$

$$\text{between subject } \widehat{\Omega} \text{ and target } \widehat{\Omega}, \quad \lambda = \arg\min_{\lambda} \sum_{a=1}^{n_{\text{subj}}} \left\|\widehat{\Omega}_a - \overline{\Omega}\right\|_2^2$$

$$\mathrm{FC}_{\text{pcorr}}(i,j) = -\frac{\widehat{\Omega}_{ij}}{\sqrt{\widehat{\Omega}_{ii}\,\widehat{\Omega}_{jj}}}, \text{3. Normalize precisionmatrix } \widehat{\Omega} \text{ to compute partial correlation FCpcorr}$$

For the Shen268 and Coco439 atlases, the target $\overline{\Omega}$ was computed by averaging the pseudoinverse of each subject FC rather than the inverse. For the Lifespan data, the regularization target was the $\overline{\Omega}$ from the HCP-YA training data. We computed $\lambda$ for each atlas based on the CRHD young adult FCcorr data, and applied that $\lambda$ the combined Lifespan dataset to generate FCpcorr. FS86 $\lambda = 0.15$, Shen268 $\lambda = 0.35$, Coco439 $\lambda = 0.58$.

**Construction of structural connectomes**

The diffusion MRI (dMRI) data were preprocessed by HCP using the minimal preprocessing pipeline, which jointly corrected for motion, susceptibility distortion and eddy current distortion using FSL's 'topup' and 'eddy' tools ([13;14], before being linearly coregistered to the anatomical image. Preprocessed dMRI were further processed using MRtrix3 (ref.[15];3.0_RC3), including bias correction, constrained spherical deconvolution (multi-shell, multi-tissue fiber orientation distribution (FOD) estimation, lmax = 8, ref.[16]), and whole-brain tractography. We performed both deterministic (sdstream[17]) and probabilistic, anatomically constrained tractography (ifod2act[18;19]) using dynamic white-matter seeding[20] for both methods, resulting in 5 million total streamlines per subject for each tractography algorithm. SC matrices were constructed for each atlas by counting the number of streamlines that ended in each pair of gray matter regions, normalized by the total volume of each region pair divided by the estimated total intracranial volume (eTIV). The eTIV normalization term is critical here as females tend to have smaller skulls and thus smaller regional volumes; if you do not normalize by eTIV the smaller regional volumes would inflate the SC entries artificially.

**Parcellations**

We used three different whole-brain atlases, covering a range in size and construction methodology. The 86-region FreeSurfer atlas (FS86) combines 68 cortical gyri from Desikan–Killiany and 18 subcortical gray matter regions (aparc+aseg output file[21;22]). The 268-region Shen atlas (Shen268) is an MNI-space cortical and subcortical volumetric atlas based on resting-state fMRI clustering[23;24]. The 439-region CocoHCP439 atlas (Coco439) combines 358 cortical regions from the HCP multimodal parcellation[25], defined using anatomical and functional connectivity gradients, with 12 subcortical regions from FreeSurfer aseg (further modified by FSL FIRST[26]; FSL 6.0), 30 thalamic nuclei derived from FreeSurfer 7.2.0 (ref.[27]; 50 original outputs included many small nuclei, which were merged into the final set of 30), 27 cerebellar regions from the SUIT atlas[28], and 12 additional subcortical nuclei from AAL3 (ref.[29]). The FS86 and Coco439 atlases were defined based on each individual subject's FreeSurfer surface and subcortical parcellations, whereas the Shen268 atlas was applied directly to MNI-resampled data for each subject. The FS86, Shen268 and Coco439 parcellations result in 3,655, 35,778 and 96,141 pairwise connectivity estimates, respectively.

### Sex-related trajectories across age

We provide an individual-level view of sex-related patterns by visualizing projections onto the sex-related latent axis (PC2) of the Krakencoder/Sex Fine-tuned Krakencoder representations as a function of age (Fig. S1). Across both models, sex separation was weakest in early childhood, increased during adolescence, and persisted into adulthood, with a slight reduction at older ages (the increased sex separation among 100+ participants is likely due to small sample size). These patterns are consistent with trends observed in classification performance and reinforce the interpretation that Krakencoder and Sex Fine-tuned Krakencoder are capable of capturing sex differentiation in brain connectivity across ages.

### Sex classification using different classifiers

Figure S2 compares sex classification performance across different models using the Sex Fine-tuned Krakencoder latent representations derived from fusion FC, fusion SC, and fusion FC and SC. Besides logistic regression, Kernel ridge regression, Support vector classifier (SVC), and Multi-layer perceptron (MLP) were used. Different classifiers had similar age-related sex patterns across lifespan but some had systematic shifts in accuracy; for example the MLP did not perform well likely due to the relatively small N per bin.

### Sex prediction for whole classification dataset

To compare the performance of the sex classification models for larger sample sizes, we summarized test accuracy across 100 independent repetitions using 50/50 train–test splits of the 838-subject sex classification dataset (Figure S3) without binning into age groups. Each model was evaluated under three input settings: Fusion (all FC and SC connectomes), Fusion FC (all FC connectomes), and Fusion SC (all SC connectomes). Bars represent the mean test accuracy across repetitions, with error bars indicating the standard deviation.

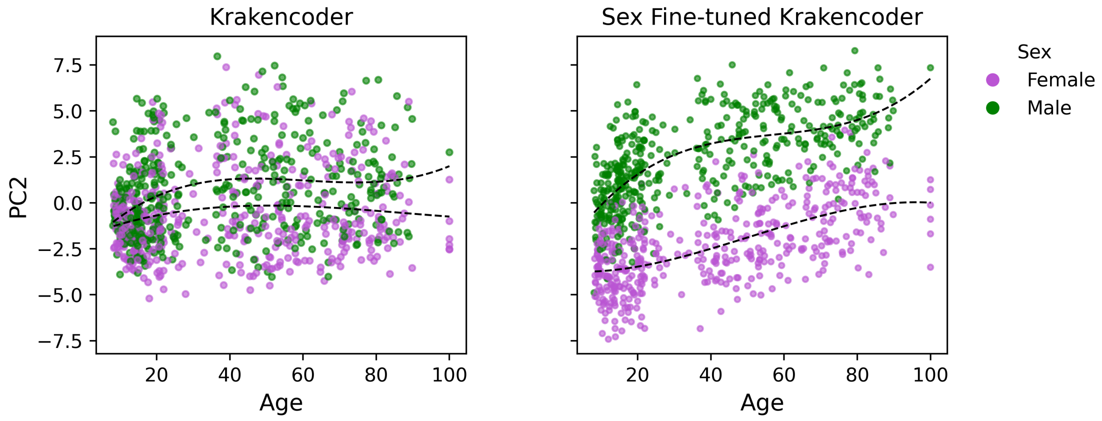


Figure S1: Individual-level visualization of sex-related patterns across the lifespan. Scatter plots show projections onto a sex-related principal component (PC2) of the latent space as a function of age for the Krakencoder (left) and the sex Fine-tuned Krakencoder (right). Each point represents an individual participant, colored by sex. Dashed curves (upper: male, lower: female) indicate sex-specific cubic smoothing splines summarizing age-dependent trends.

Overall, when ample training data are available, all three models achieve high accuracy in sex prediction, with Sex fine-tuned Krakencoder slightly outperforming the other models. Notably, the main results indicate that the Sex fine-tuned Krakencoder exhibits greater robustness at smaller sample sizes than the ensemble approach, while also offering improved computational efficiency.

### Mean framewise displacement

We provide violin plots to visualize and compare the distributions of mean framewise displacement (FD) during the fMRI scans across age and sex groups, see Figure S4 Panel (b). The distribution of mean FD centered at 0.1-0.2mm across all ages, with younger children (ages 8–11) and older adults (ages 65+) showing higher levels of head motion compared to adolescents and young adults. Males and females did not have significantly different mean FD in any age group, all t-tests had corrected $p > 0.05$.

### Site effects

When analyzing sex differences in FCs or SCs across age groups, data are often collected from multiple imaging centers or scanners at different sites. Each site may introduce systematic variations due to differences in hardware, acquisition protocols, preprocessing pipelines, or participant recruitment strategies. If each age bin contains subjects with uneven site by sex distributions, observed differences in connectivity between the sexes could be confounded by site-specific effects. This problem becomes critical in lifespan analyses, where age-related trends in sex differences may be distorted if site proportions vary systematically across bins.

We applied harmonization procedures to mitigate site effects and confirmed that multi-

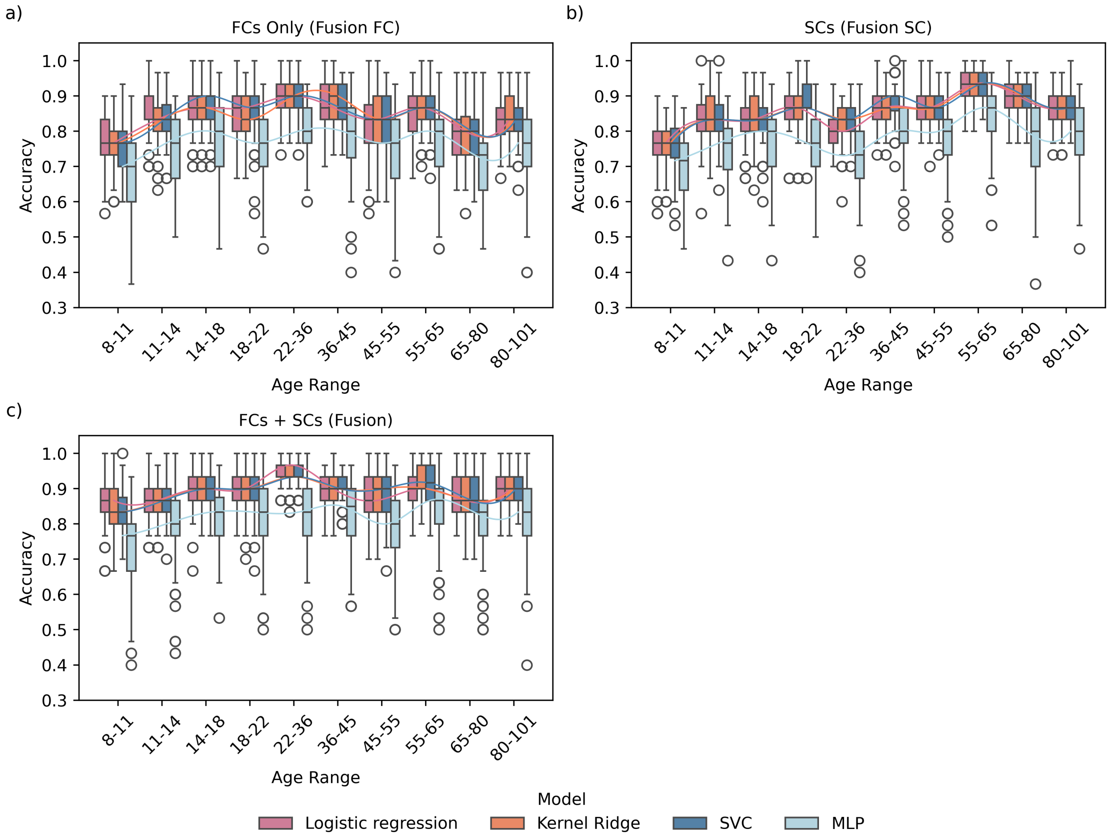


Figure S2: Sex classification accuracy built on Sex fine-tuned Krakencoder latent representations using different classifiers. Boxplots show test accuracy from 100 repetitions for four models (logistic regression, kernel ridge regression, SVC, and MLP). Panels compare latent spaces learned from (a) Fusion FC, (b) Fusion SC, and (c) Fusion (FC+SC). For each model, a spline was fit to the average accuracy to visualize age-related trends in sex classification accuracy.

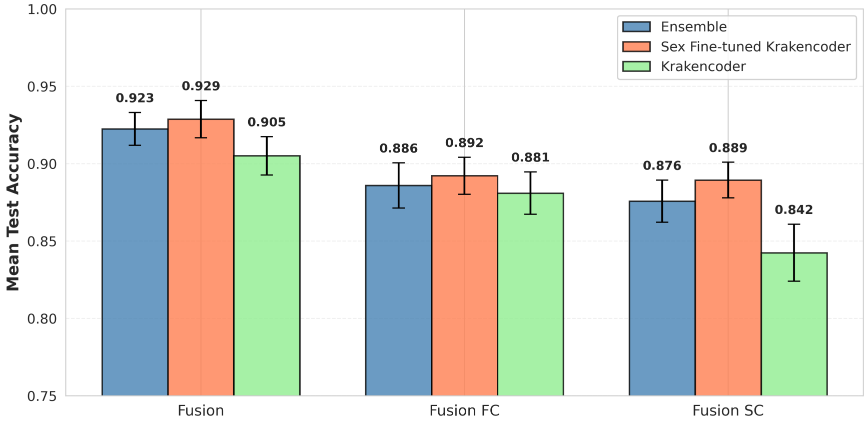


Figure S3: Sex prediction for whole classification dataset (100 repetitions, 50/50 train/test)

site data collection had minimal influence on our sex difference analysis. Figure S4 (a) shows the distribution of sites across age bins, with the first bar of each age bin representing females and the second representing males. Between males and females in the same age range, the site proportions are similar. Across age, the site proportions are similar except for age bin 22-36, and one site (Harvard) with samples only in the HCP-D range and another site (MGH) having samples only in the HCP-A range. We further fine-tuned Krakencoder (*Sex - Site Fine-tuned Krakencoder*) by incorporating a penalty on site classification, and applied CovBat[30] to the raw connectome data for the ensemble technique (*Ensemble Covbat*). The resulting sex prediction accuracies (Figure S5 (b)) show consistent patterns with those observed before removing site effects, indicating that they are not largely influential in the connectome data. Figure S5 (c) illustrates site prediction accuracy, which remains generally low across bins (with a modest increase at ages 22–36), indicating that site effects are not a dominant factor in our dataset. Notably, both the *Ensemble CovBat* and *Sex–Site Fine-tuned Krakencoder* achieve substantially lower site prediction accuracy in only the 22–36 group compared with the baseline *Ensemble*, *Krakencoder*, and *Sex Fine-tuned Krakencoder* models across all fusion modalities, confirming that site effects were minimal. Together, these results demonstrate that site does not substantially influence our analysis.

### Supplementary Results for Sensitivity Analysis

Figures S6 and S7 provide the details for Figure 2.

### Ensemble Model Feature Importance

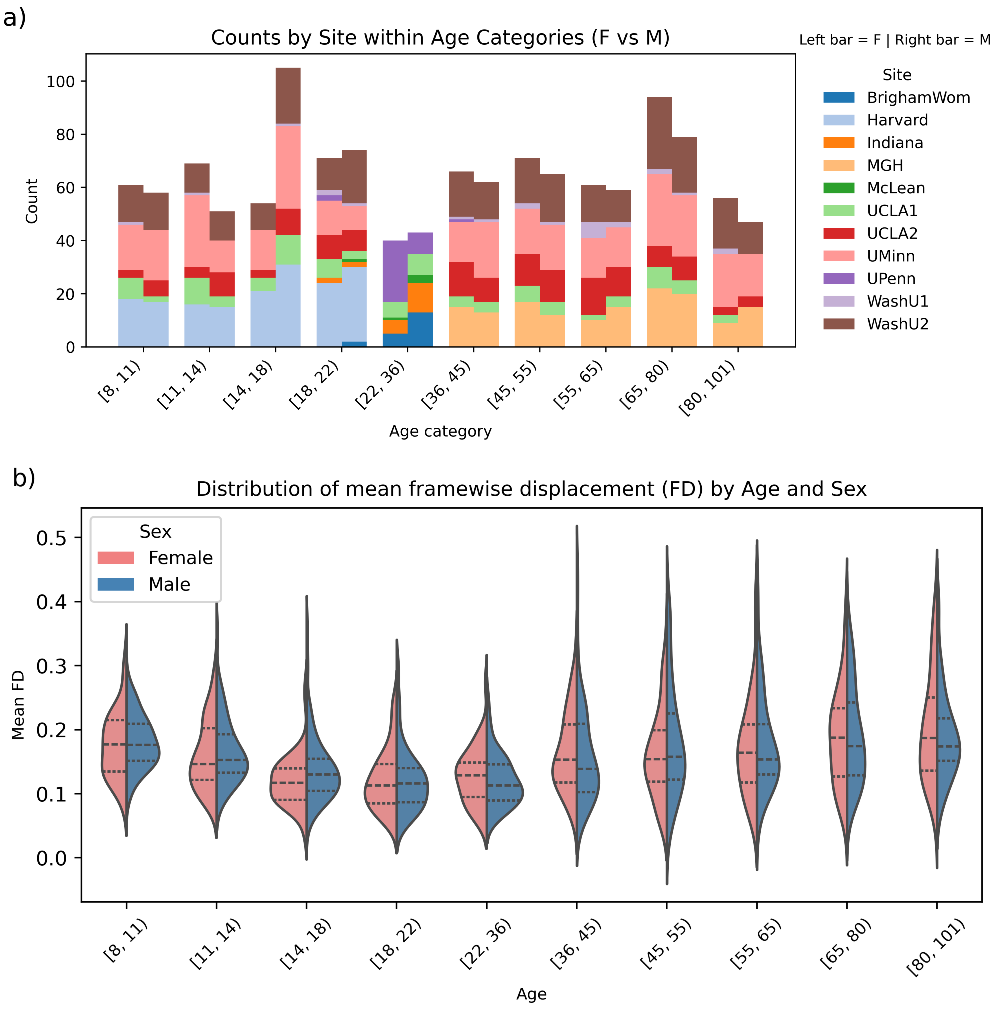


Figure S4: Panel (a): Distribution of site counts across age bins, separated by sex. Each age bin shows two bars, where the first bar are females (F) and the second bar are males (M). Pnael (b): Violin plots of mean framewise displacement (FD) during the fMRI scan across age categories, separated by sex (female: red, male: blue). Each violin shows the distribution within an age bin, with quartile markers indicated inside the split violins.

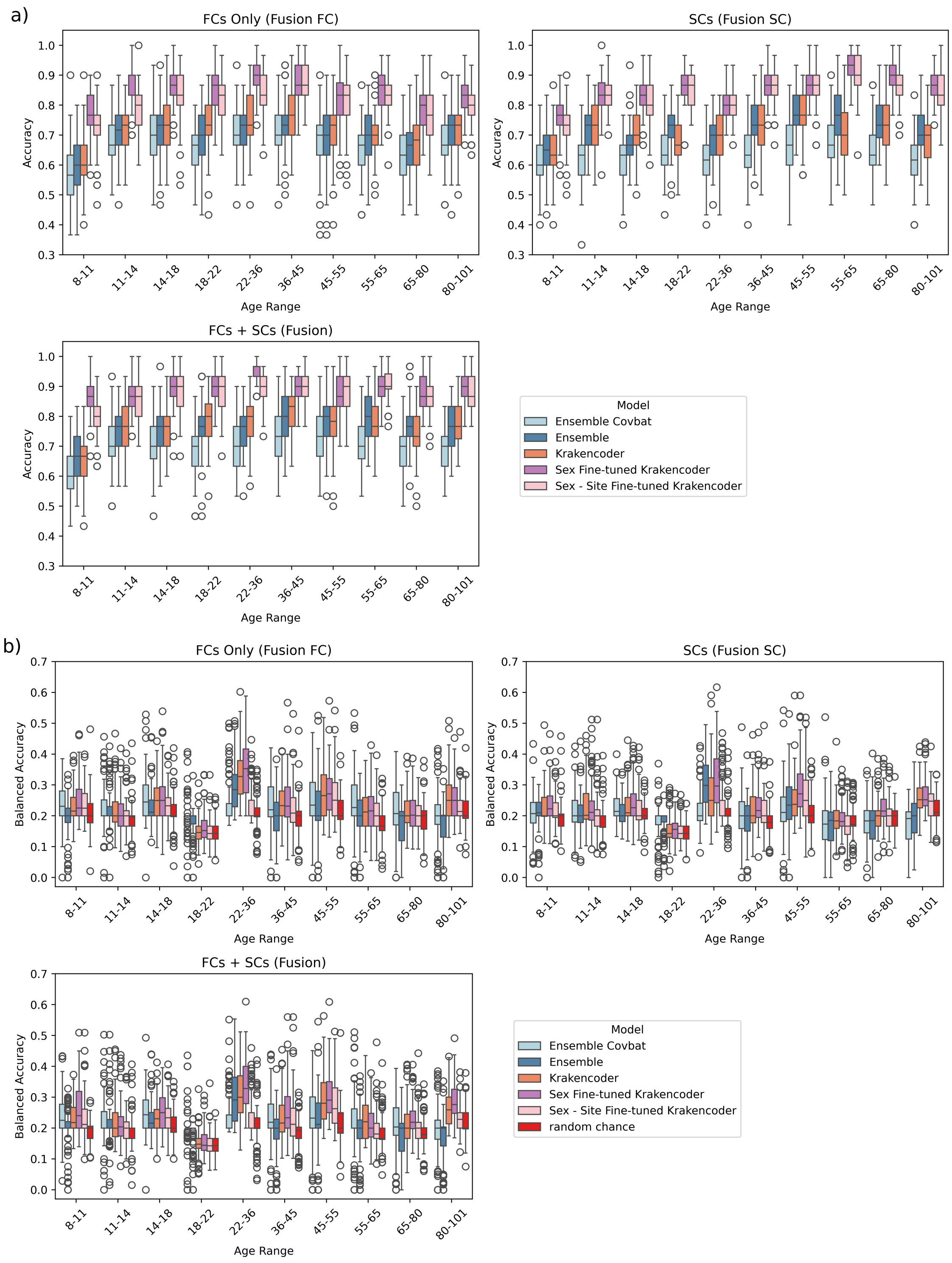


Figure S5: Panel (a): Distribution of site counts across age bins, separated by sex. Each age bin shows two bars, where the first bar are females (F) and the second bar are males (M). Panel (a): Sex classification accuracies across the lifespan. Panel (b): Site classification balanced accuracies across the lifespan, with red bars indicating random chance balanced accuracy obtained by permuting site labels and generating predictions using the sex and site fine-tuned Krakencoder.

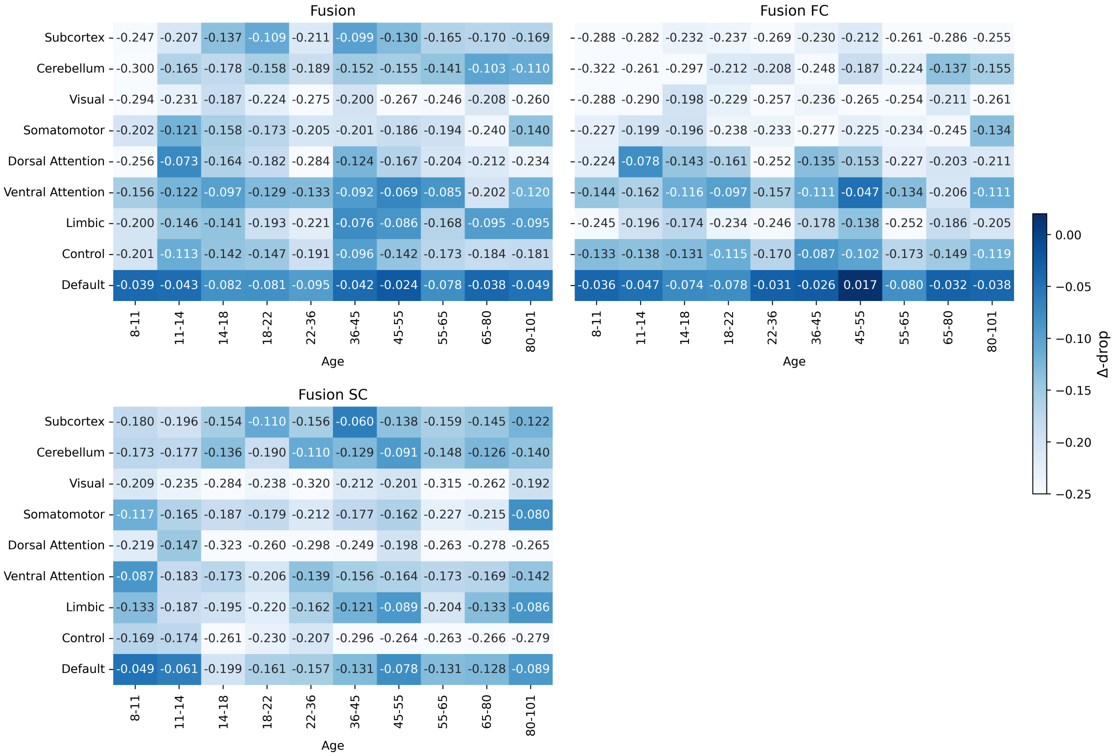


Figure S6: Heatmaps of average sex-classification accuracy drop (Δ-drop) across age bins using the sex fine-tuned Krakencoder model. Each cell shows the mean Δ-drop value, with darker blue indicating smaller decreases in sex-classification accuracy, indicating stronger sex association.

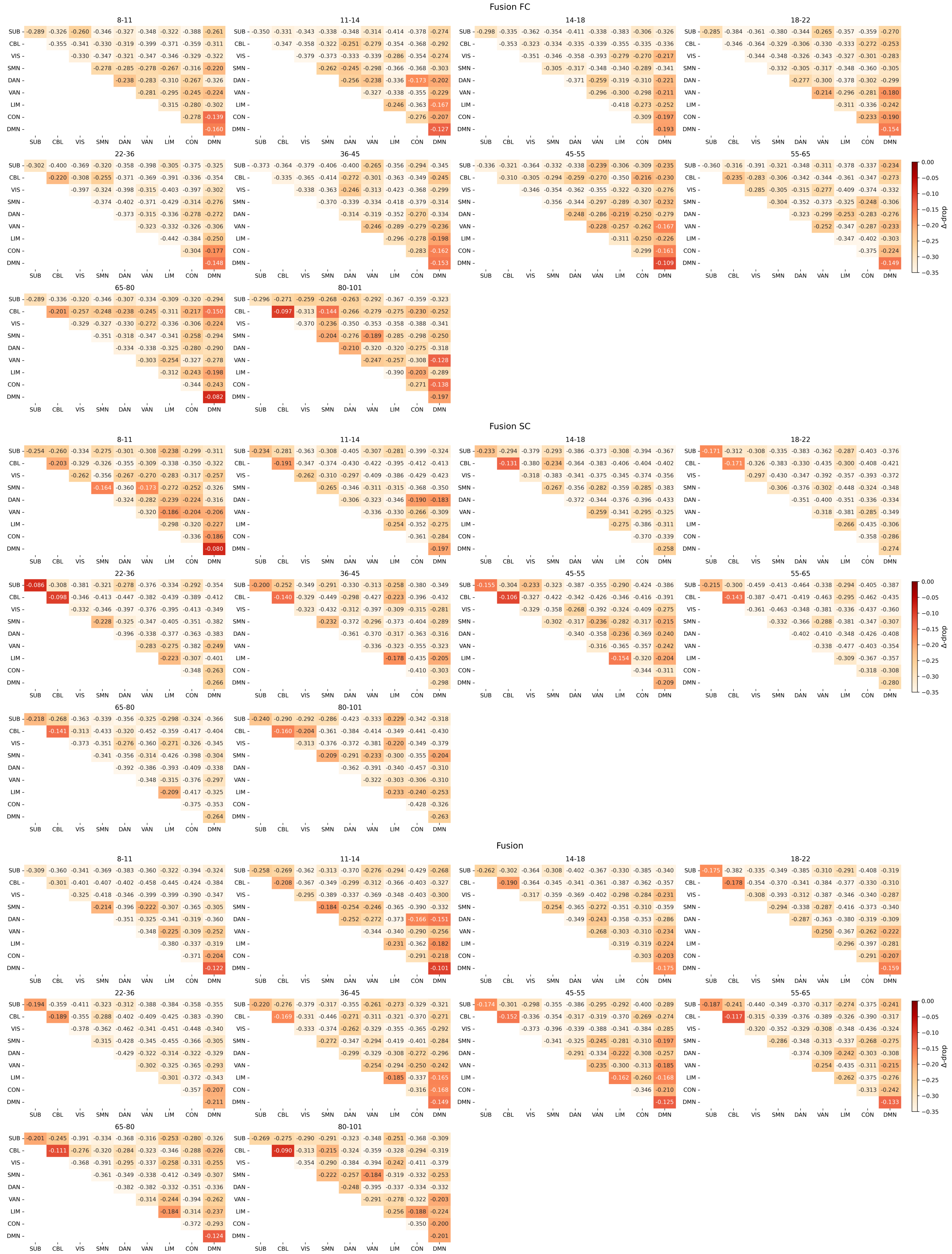


Figure S7: Mean sex-classification accuracy drops (Δ-drops) when connections between those pairs of networks were retained using the sex fine-tuned Krakencoder model. Darker red indicates smaller decreases in sex classification accuracy, indicating stronger sex association.

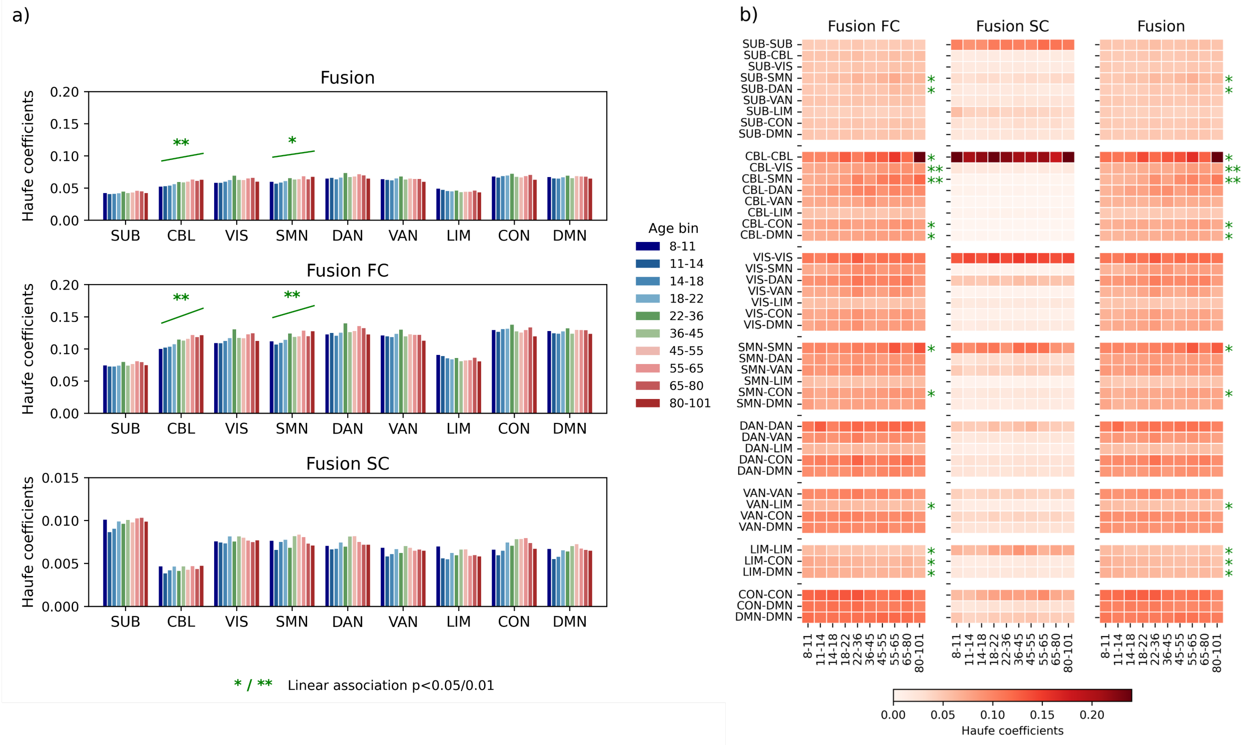


Figure S8: Panel (a): Network-level average feature importances from the ensemble model. For each age bin, the absolute values of the Haufe coefficients were averaged across all connections to and from each network to quantify overall importance. Panel (b): Network-pair average feature importances from the ensemble model. For each age bin, the absolute values of the Haufe coefficients were averaged across all connections within each network pair to quantify pairwise feature importance. Solid curves indicate significant linear age trends; no quadratic trends were statistically significant. P-values were corrected using FDR-control across the 9 networks within each modality (Fusion FC and Fusion SC) by FDR control.

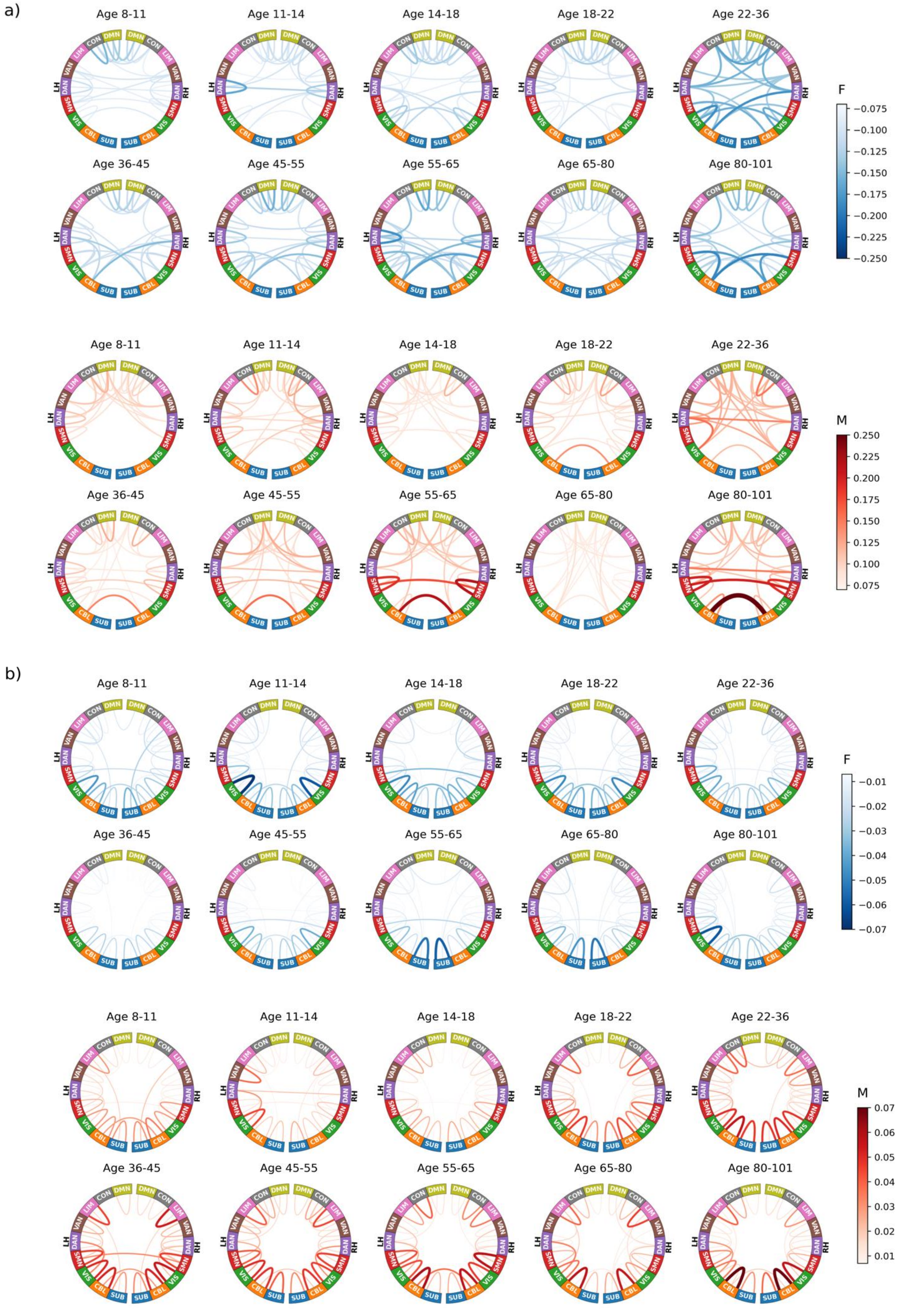


Figure S9: Sex-specific feature importance patterns for FC (a) and SC (b). Top 15% strongest connections are shown for each age.

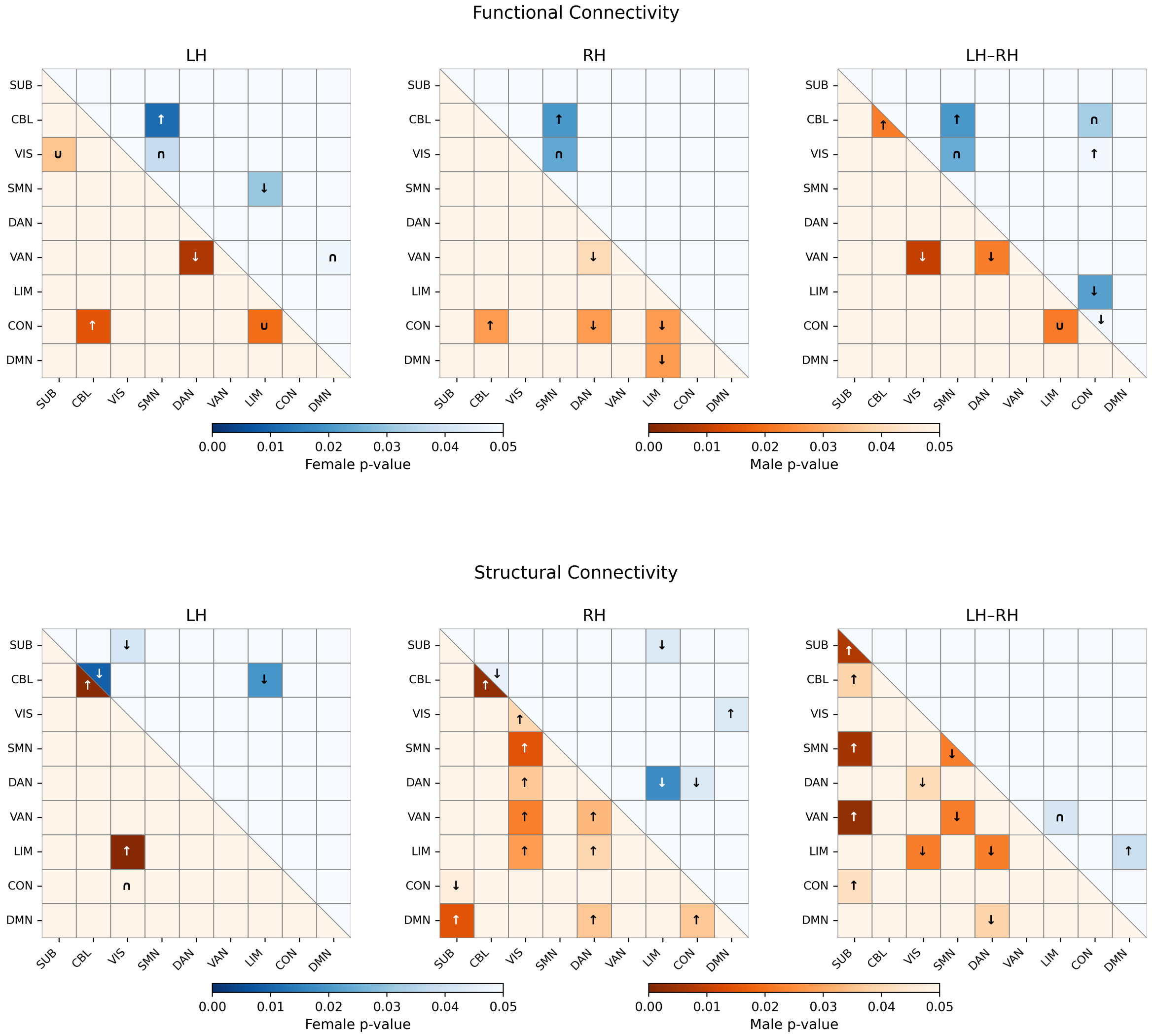


Figure S10: Network-pair–level significance of linear and quadratic age trends in FC (Top) and SC (bottom) for Figures 3 and 4. Each $9 \times 9$ matrix shows results for the left hemisphere (LH), right hemisphere (RH), and inter-hemispheric (LH–RH) connections. The upper triangle represents females and the lower triangle represents males. Color intensity encodes the minimum p-value across linear (Spearman) and quadratic tests (darker = more significant). Overlaid arrows (↑ / ↓) indicate significant (corrected $p < 0.05$) linear increases or decreases with age, and U- or inverted-U-shaped symbols denote the significant quadratic associations. Color bars indicate p-value scales separately for males (orange) and females (blue). P-values were adjusted using FDR control, applied separately within each hemisphere and modality (Fusion FC or Fusion SC) across all network pairs.

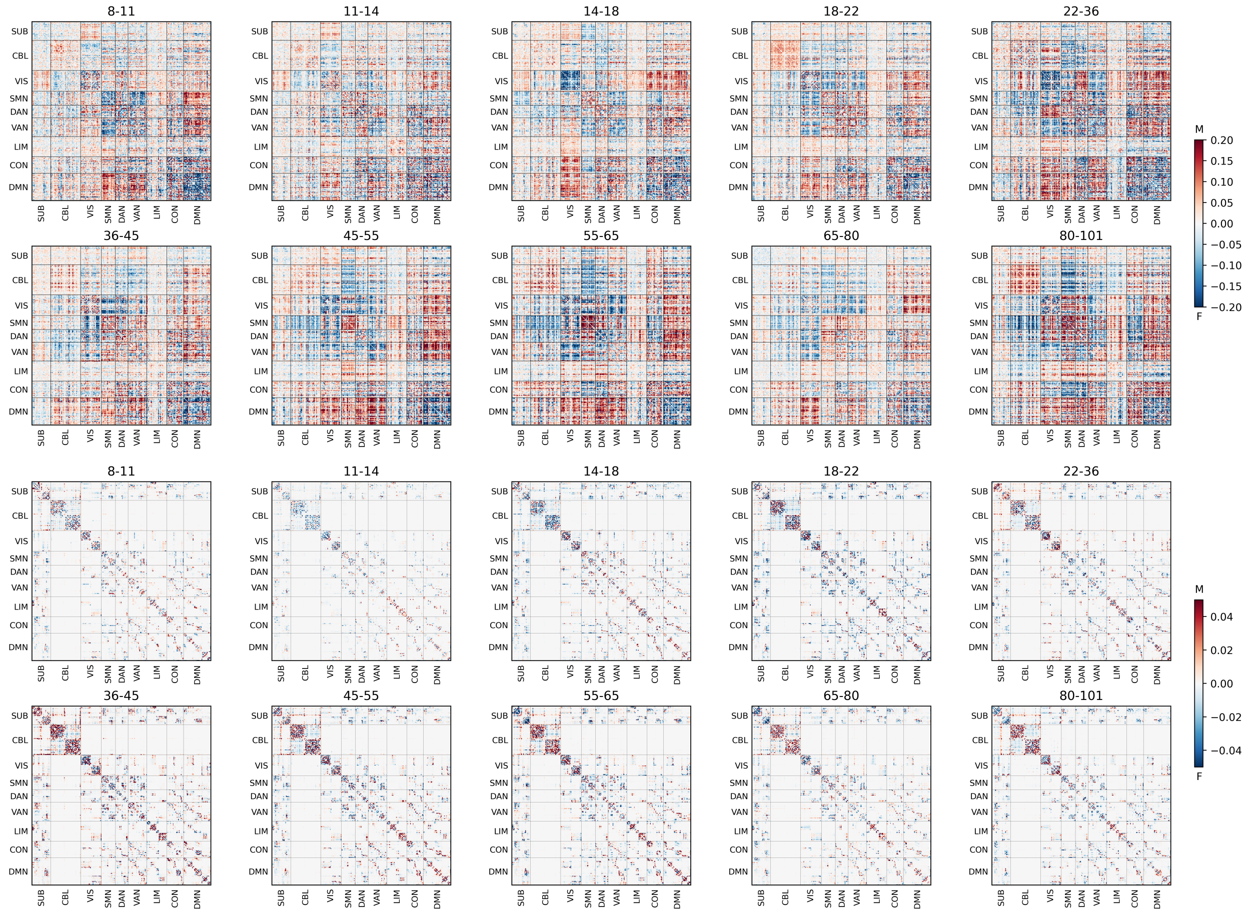


Figure S11: The Heatmaps show feature importance (averaged standardized Haufe coefficients) where more blue indicates stronger connections are related to female sex and red indicates stronger connections are linked to male sex. Top two rows: FC using Pearson correlation, the shen268 atlas, high-pass filtering, and global signal regression; Bottom two rows: SC using the shen268 atlas, probabilistic tractography (ifod2), and anatomical constraint global filtering (act).. Regions are ordered by functional Yeo networks, with grey lines marking network boundaries.
